# Multivalent GU-rich oligonucleotides sequester TDP-43 in the nucleus by inducing high molecular weight RNP complexes

**DOI:** 10.1101/2023.08.01.551528

**Authors:** Xi Zhang, Tanuza Das, Tiffany F. Chao, Vickie Trinh, Rogger Carmen, Jonathan Ling, Petr Kalab, Lindsey R. Hayes

## Abstract

TDP-43 nuclear clearance and cytoplasmic aggregation are hallmarks of TDP-43 proteinopathies. We recently demonstrated that binding to endogenous nuclear GU-rich RNAs sequesters TDP-43 in the nucleus by restricting its passive nuclear export. Here, we tested the feasibility of synthetic RNA oligonucleotide-mediated augmentation of TDP-43 nuclear localization. Using biochemical assays, we compared the ability of GU-rich oligonucleotides to engage in multivalent, RRM-dependent binding with TDP-43. When transfected into cells, (GU)16 attenuated TDP-43 mislocalization induced by transcriptional blockade or RanGAP1 ablation. Clip34nt and (GU)16 accelerated TDP-43 nuclear re-import after cytoplasmic mislocalization. RNA pulldowns confirmed that multivalent GU-oligonucleotides induced high molecular weight RNP complexes, incorporating TDP-43 and possibly other GU-binding proteins. Transfected GU-repeat oligos disrupted TDP-43 cryptic exon repression, likely by diverting TDP-43 from endogenous RNAs, except for Clip34nt which contains interspersed A and C. Thus, exogenous multivalent GU-RNAs can promote TDP-43 nuclear localization, though pure GU-repeat motifs impair TDP-43 function.

## INTRODUCTION

TDP-43 is an essential nucleic-acid binding protein that regulates many aspects of RNA metabolism, including pre-mRNA splicing and stability.^1^ Like other heterogeneous nuclear ribonucleoproteins (hnRNPs), TDP-43 shuttles between the nucleus and cytoplasm and is predominantly nuclear in healthy cells.^2,3^ In ‘TDP-43 proteinopathies’, a class of neurodegenerative disorders that includes amyotrophic lateral sclerosis (ALS), frontotemporal dementia (FTD), limbic-predominant age-related TDP-43 encephalopathy (LATE), and related disorders, affected cells in the central nervous system show TDP-43 nuclear clearance and cytoplasmic aggregation.^4^ Though TDP-43 mutations have been identified in rare genetic cases,^5^ most patients with TDP- 43 proteinopathy have sporadic, non-inherited disease. Despite advances in understanding the regulation of TDP-43 nucleocytoplasmic transport and the downstream consequences of TDP-43 mislocalization,^6^ the molecular events initiating TDP-43 mislocalization in neurodegeneration remain obscure. The development of therapies targeting TDP-43 nuclear clearance, loss of function, and cytoplasmic aggregation is of major interest toward slowing the progression of neurodegeneration.^7^

TDP-43 is a 43-kDa protein with an N-terminal oligomerization domain followed by a nuclear localization sequence (NLS), tandem RNA recognition motifs (RRM1 and RRM2), and a C-terminal intrinsically disordered region that mediates liquid-liquid phase separation (LLPS).^8^ Structural analysis demonstrates that the RNA binding groove of RRM1-2 contacts 10 nucleotides of a 5′-GNGUGNNUGN-3′ motif, with ‘AUG12’ (5’- GUGUGAAUGAAU-3’) binding TDP-43 stably in a single conformation.^9^ Crosslinking immunoprecipitation (CLIPseq) from cell lines, mouse, and postmortem human brain confirms that TDP-43 binds GU- and G,U,A-rich RNA motifs primarily within introns, as well as a subset of noncoding RNAs, miRNAs, and mRNAs.^10–12^ Notably, TDP-43 binds its own 3’-UTR at a sequence termed Clip34nt to autoregulate its expression.^10,13^ Binding up to one-third of the mammalian transcriptome, TDP-43 functions prolifically in the regulation of RNA processing, including alternative splicing,^10,11^ the repression of cryptic exons,^12,14,15^ alternative polyadenylation,^16–20^ and the processing of miRNA^21^ and lncRNAs, including long interspersed nuclear elements (LINEs).^22^

The nucleocytoplasmic localization of TDP-43 is regulated by a dynamic balance between nuclear import and export. The active nuclear import of TDP-43 primarily occurs via NLS binding to importins α and β.^2,23^ In addition, the recently discovered binding of importin β to the C-terminal domain of TDP-43 revealed a potential NLS-independent import pathway for TDP-43-ΔNLS.^24^ Because TDP-43-ΔNLS showed only limited nuclear entry relative to wild-type TDP-43 in heterokaryon assays,^2^ the NLS-independent import may be significantly slower. The nuclear export of full-length TDP-43, initially thought to occur via exportin-1 (XPO1) binding to a putative nuclear export signal (NES) in the RRM2 domain, was subsequently shown to occur by passive diffusion through nuclear pore channels and is exquisitely size-dependent.^25–28^ In multiple export assays, the addition of bulky tags markedly delayed TDP-43 nuclear export by raising its size above the permeability barrier of the nuclear pore complex (NPC), which increasingly excludes cargoes above 30-60 kDa in size.^29^ An exception is the short TDP-43 isoform, which is upregulated by pathologic hyperexcitability and localizes to the cytoplasm via a distinct NES for XPO1 arising in the C-terminus.^30^

We recently demonstrated that the nuclear localization of full-length TDP-43 critically depends on binding to nuclear RNAs, which sequester TDP-43 in the nucleus and restrict its availability for passive nuclear export.^28^ In permeabilized and live cell nuclear export assays, depletion of nuclear RNA binding sites for TDP-43, via RNase degradation or inhibition of transcription, released TDP-43 for passive diffusion from the nucleus. RRM mutations or displacement of TDP-43 from endogenous nuclear RNAs with monovalent (GU)6 oligonucleotides also promoted cytoplasmic mislocalization. Conversely, inhibiting splicing to promote the accumulation of nuclear intronic binding sites for TDP-43 promoted TDP-43 nuclear retention. Based on these data, we hypothesized that longer exogenous GU-rich oligonucleotides might augment TDP-43 nuclear localization, provided the oligonucleotides form multivalent complexes with TDP-43 large enough to restrict its diffusion through the NPC.

Here, we report the characterization and functional outcome of GU-rich oligonucleotides tested for their ability to promote TDP-43 nuclear retention. The TDP-43 binding valency of GU- and G,U,A-rich oligonucleotides with ≥2 predicted TDP-43 binding sites, of differing lengths and sequences, was tested in biochemical assays and living cells. (GU)16 > Clip34nt > (‘AUG12’)2 induced TDP-43 RRM1,2 multimerization *in vitro* and promoted the formation of heterogenous, high molecular weight TDP-43 complexes in living cells. Remarkably, (GU)16 attenuated TDP-43 mislocalization following transcriptional blockade and RanGAP1 ablation, confirming that exogenous oligonucleotides can scaffold TDP-43 and promote its nuclear retention. Clip34nt and (GU)16 also accelerated TDP-43 nuclear re-import after cytoplasmic mislocalization. However, transfected oligos containing tandem GU-repeats caused unwanted disruption of TDP-43 cryptic exon repression, likely by diverting TDP-43 from endogenous RNAs. Clip34nt, perhaps due to its lower-affinity, A,C-interspersed sequence, largely spared TDP-43 function both by a cryptic exon reporter and at endogenous cryptic targets, indicating that further optimization of oligo motifs may enable RNA-mediated promotion of TDP-43 nuclear localization without compromising its function.

## RESULTS

### Multivalent interactions of GU-rich oligonucleotides with TDP-43

GU-rich oligonucleotide sequences were selected with well-characterized TDP-43 binding motifs and the established or predicted ability to form multimers with TDP-43 (fig 1A). The multivalent binding of TDP-43 to GU-rich RNAs is well demonstrated. Endogenous TDP-43 RNA binding sites frequently occur in clusters that coordinate multivalent binding depending on sequence, length, and density.^31^ Biochemical assays confirm that the valency of TDP-43 binding to synthetic (GU)n oligonucleotides is length-dependent, with monovalent binding ≤(GU)6 and multivalent binding ≥(GU)12.^32^ Pure GU-repeats and the ‘AUG12’ motif exhibit low nanomolar affinity (0.8-26.9 nM) for TDP-43 in *in vitro* assays,^9,33–35^ while Clip34nt affinity has been estimated at 97-520 nM.^34,36^ Clip34nt is a 34-base G, U, and A-rich sequence that strongly promotes TDP-43 LLPS^33^ and has shown promise as a chaperone to disaggregate cytoplasmic TDP-43.^37^ The Clip34nt binding valency for TDP-43 has not yet been established.

**Figure 1.**
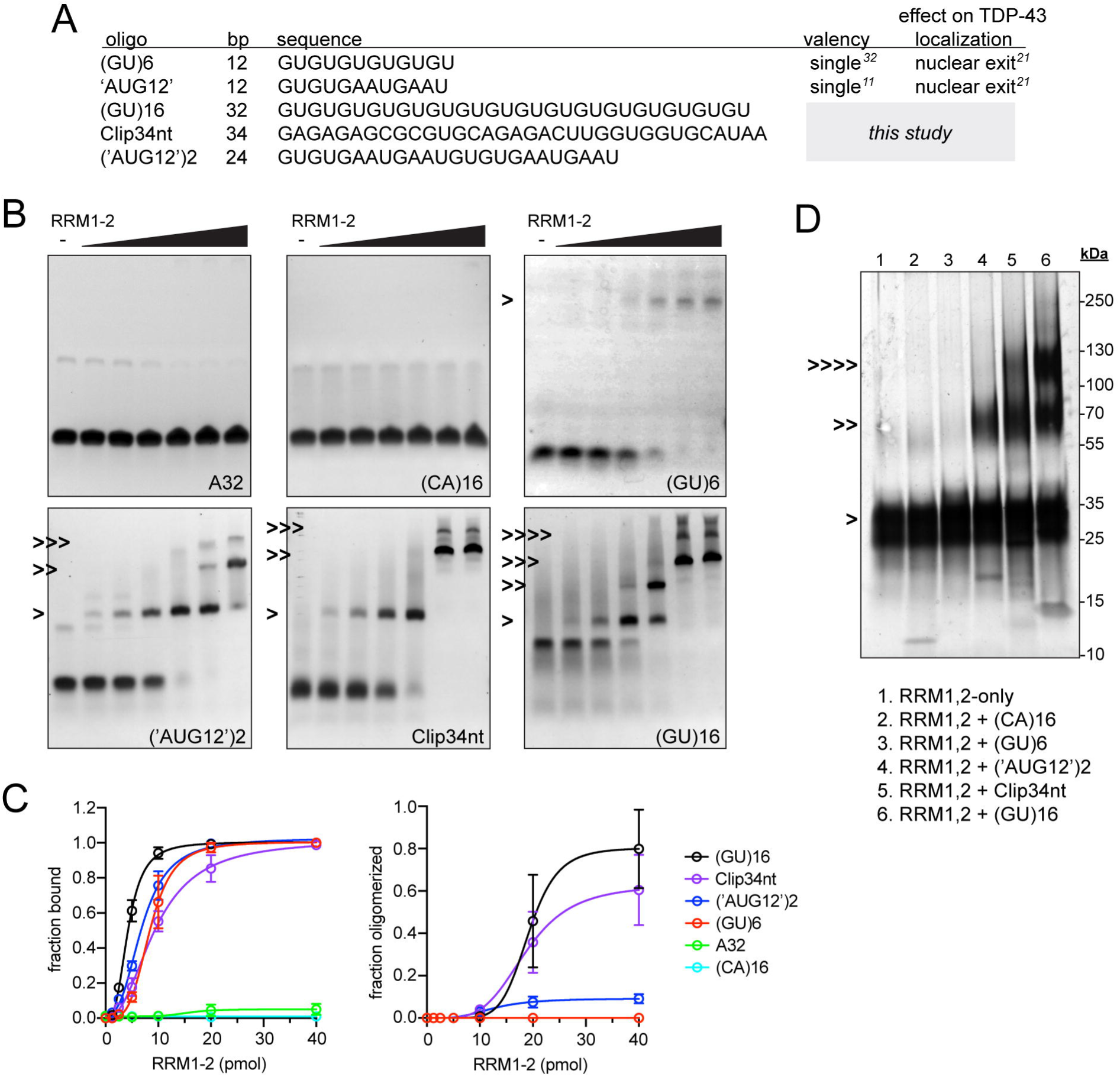
GU-rich oligonucleotides bind TDP-43 RRM1,2 with variable multivalency. A. Oligonucleotide sequences tested in this study. For base modifications see **table S1.** B. EMSA assays in which 10 pmol oligonucleotide was mixed with 0-40 pmol recombinant RRM1-2 protein, resolved under native conditions, and visualized with SYBR gold nucleic acid stain. Band shifts corresponding to predicted monomers (>), dimers (>>), trimers (>>>), or tetramers (>>>>) are indicated. C. Left: total bound RNA fraction from EMSA assays in (B), quantified as the sum of all shifted bands divided by the total RNA signal. Right: oligomerized RNA fraction from EMSA assays in (B), quantified as the sum of bands predicted to contain >2 RRM1,2 molecules, divided by the total RNA signal. Mean ± SD is shown for 3-6 biological replicates. D. BS3-crosslinked RRM1,2-oligonucleotide complexes (10 pmol oligo + 40 pmol protein) analyzed by SDS- PAGE and visualized with silver stain. Bands corresponding to predicted monomers (>), dimers (>>), and tetramers (>>>>) are indicated. Representative of 3 biological replicates.

(GU)16, (‘AUG12’)2, and Clip34nt oligonucleotides were synthesized with 2’-O-methyl and phosphorothioate (PS)-modifications commonly utilized to stabilize antisense oligonucleotides (ASO) against RNase degradation^38^ and binding to recombinant SUMO-tagged RRM1,2 was analyzed. First, electrophoretic mobility shift assays (EMSAs) were performed by combining each oligonucleotide with increasing concentrations of RRM1,2 and RNA-protein complexes were resolved by native electrophoresis (fig 1B-C). All three GU-rich oligonucleotides readily bound RRM1,2. A32 and (CA)16 were utilized as controls for specificity, and no appreciable binding with RRM1,2 was observed. As previously reported,^32^ (GU)6 showed a single band shift with increasing RRM1,2 concentrations, indicating strictly monovalent complexes. (‘AUG12’)2, Clip34nt, and (GU)16 all exhibited multiple, discrete band shifts with increasing protein concentrations, suggesting multimers. Quantifying band shifts likely to involve more than two RRM1,2 molecules showed that (GU)16 had the highest propensity for multivalency, followed by Clip34nt (fig 1C). Band shifts corresponding to complexes with >2 RRM1,2 were rare for (‘AUG12’)2. To determine the molecular weight of the RNA-protein complexes, the amine-crosslinker bis(sulfosuccinimidyl)suberate (BS3) was utilized before SDS-PAGE and proteins were visualized with silver stain (fig 1D). As predicted from the EMSAs, (GU)6 formed monomers, (‘AUG12’)2 formed a mix of monomers and dimers, and Clip34nt and (GU)16 additionally formed ∼120-130 kDa complexes, likely corresponding to tetramers of RRM1,2. The putative tetrameric complexes appeared most abundant with (GU)16 although silver stain is poorly quantitative.

### Binding valency predicts the effect of GU-rich oligonucleotides on TDP-43 nuclear localization

Previously, we showed that (GU)6 rapidly entered transfected HeLa cell nuclei, bound TDP-43, and promoted nuclear efflux, likely by competitive displacement of TDP-43 from endogenous nuclear RNAs, freeing the resulting ∼45 kD complex for passive diffusion through nuclear pore channels.^28^ Since (GU)6 exclusively forms monovalent complexes with TDP-43 (fig 1B-C), next we tested multivalent GU-oligonucleotides along with a (CA)16 control. A 3’-biotin label was utilized to enable oligonucleotide visualization and biochemical analysis. 5 h post-transfection, cells were fixed and labeled with fluorescently tagged streptavidin (SAV). (CA)16-bio, (‘AUG12’)2-bio, Clip34nt-bio, and (GU)16-bio readily entered transfected cells (fig 2A). The N/C partitioning of the oligonucleotides differed, with (CA)16 and (‘AUG12’)2 showing nuclear enrichment and Clip34nt and (GU)16-bio showing more equivalent nuclear and cytoplasmic signal (fig 2B). TDP-43 immunolabeling demonstrated that (‘AUG12’)2, which showed the lowest propensity for multivalent binding, elicited dose-dependent TDP-43 nuclear exit (fig 2C-D), reminiscent of (GU)6.^28^ Clip34nt elicited modest nuclear exit. In contrast, (GU)16 did not alter the steady-state localization of TDP-43. Thus, the tendency of GU-rich oligonucleotides to elicit TDP-43 nuclear efflux inversely correlated with TDP-43 binding valency. Interestingly, (CA)16, which showed the strongest nuclear localization (fig 2B), increased the TDP-43 N/C ratio. The absence of direct interactions with RRM1,2 in EMSA (fig 1B,C) and with endogenous TDP-43 in RNA pulldowns (**fig 4A**) indicate that (CA)16 affected TDP-43 nuclear localization through an indirect mechanism.

**Figure 2.**
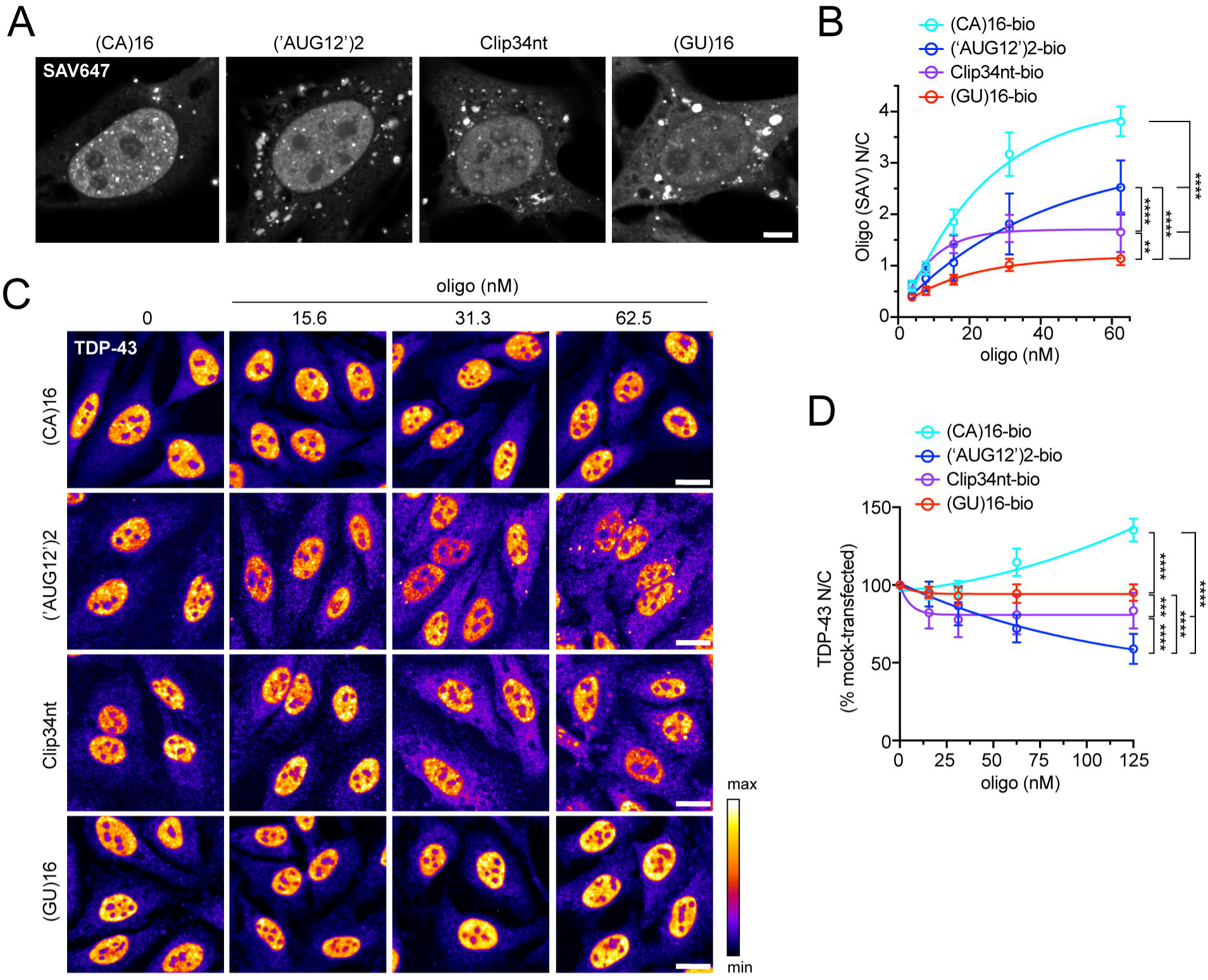
GU-oligonucleotide binding valency predicts the effect on TDP-43 localization in living cells. A. Streptavidin (SAV)-647 fluorescence in HeLa cells 5 h-post transfection with biotinylated oligonucleotides (62.5 nM). Scale bar = 5 µm. B. SAV-647 N/C 5 h post-transfection. Mean ± SD of ≥4 biological replicates. NS= not significant, ***p<0.001, ****p<0.0001 by 2-way ANOVA with Tukey’s post-hoc test. C. TDP-43 immunofluorescence in oligonucleotide-transfected HeLa cells at 5 h. The intensity histogram for each image was independently spread between the dimmest and brightest pixels and a pseudo-color linear LUT covering the full range of the data was applied (see legend). Scale bar = 20 µm. See also **figure S1**. D. TDP-43 N/C (% untreated) in oligonucleotide-transfected HeLa cells at 5 h. Mean ± SD of ≥8 biological replicates. ***p<0.001, ****p<0.0001 by 2-way ANOVA with Tukey’s post-hoc test.

Beyond shifts in nuclear/cytoplasmic localization, oligonucleotide-transfected cells displayed variable changes in the TDP-43 distribution pattern. SAV labeling showed that, in addition to diffuse signal, all four oligonucleotides formed scattered puncta in the cytoplasm suggesting a tendency to demix and self-associate. In (‘AUG12’)2 and Clip34nt-transfected cells, but not in (GU)16- or (CA)16-transfected cells, cytoplasmic TDP- 43 puncta were also observed by 5 h post-transfection (fig 2C). Next, we investigated oligonucleotide-induced alterations of nuclear TDP-43 granules (**fig S1**), predicting that the formation of TDP-43-oligo RNP complexes may alter TDP-43 granule morphology. Although no apparent alterations were seen in HeLa nuclei at 20x resolution (not shown), we tested U2OS cells where we have observed that endogenous TDP-43 nuclear RNP granules are readily visualized in untreated cells (**fig S1A**). By 24 h, U2OS cells transfected with GU oligonucleotides, particularly with (GU)16 and (‘AUG12’)2, showed a reduction in the largest nuclear granules detectable at 20x (≥0.5 µm^2^, **fig S1B-E**). To detect the suspected GU-oligo-induced increase in small nuclear granules, we imaged a subset of cells at 63X/Airyscan resolution. Indeed, (GU)16 induced a significant increase in small nuclear granules (mean area ∼0.1 µm^2^), which doubled in number compared to mock-transfected or (CA)16-transfected cells (**fig S1F-H**). Presumably, the formation of relatively small RNPs composed of high-affinity, multivalent TDP-43-(GU)16 complexes might be involved in both the reduction in nuclear TDP-43 granule size in U2OS cells (**fig S1F-H**), and in the paucity of cytoplasmic TDP-43 foci in (GU)16-transfected HeLa cells (**fig. 2C**). However, the mechanism(s) responsible for these observations await future investigation.

### Multivalent GU-oligonucleotides promote TDP-43 nuclear localization

Since (GU)16 showed the highest potential for multivalent TDP-43 interactions (**fig 1**) and did not induce TDP- 43 nuclear efflux (**fig 2**), we predicted that the size of nuclear (GU)16-TDP-43 complexes may exceed the passive diffusion limits of nuclear pore channels. To test that hypothesis, (GU)16-transfected cells were treated with the RNA Pol II inhibitor NVP2 to block transcription and deplete the nucleus of TDP-43 binding sites within nascent pre-mRNAs. As we previously reported,^28^ NVP2 rapidly induced TDP-43 cytoplasmic mislocalization in untransfected cells (fig 3A-C). Remarkably, NVP2-induced TDP-43 mislocalization was attenuated in (GU)16- transfected cells in a dose-dependent manner. At 62.5 nM (GU)16, NVP2-induced TDP-43 nuclear efflux was completely abolished. Specifically, (GU)16 attenuated the decrease in nuclear intensity and increase in cytoplasmic intensity induced by NVP2, confirming the blockade of TDP-43 translocation to the cytoplasm (**fig S2A**). In contrast, NVP2-induced TDP-43 cytoplasmic mislocalization was essentially unaltered in Clip34nt and (‘AUG12’)2 transfected cells (fig 3D-E). Unexpectedly, though consistent with its effect in non-NVP2 treated cells (fig 2C-D), 62.5 nM (CA)16 partially attenuated NVP2-induced TDP-43 mislocalization (fig 3F). Further investigation of the indirect mechanism for (CA)16-induced TDP-43 nuclear localization is ongoing.

**Figure 3.**
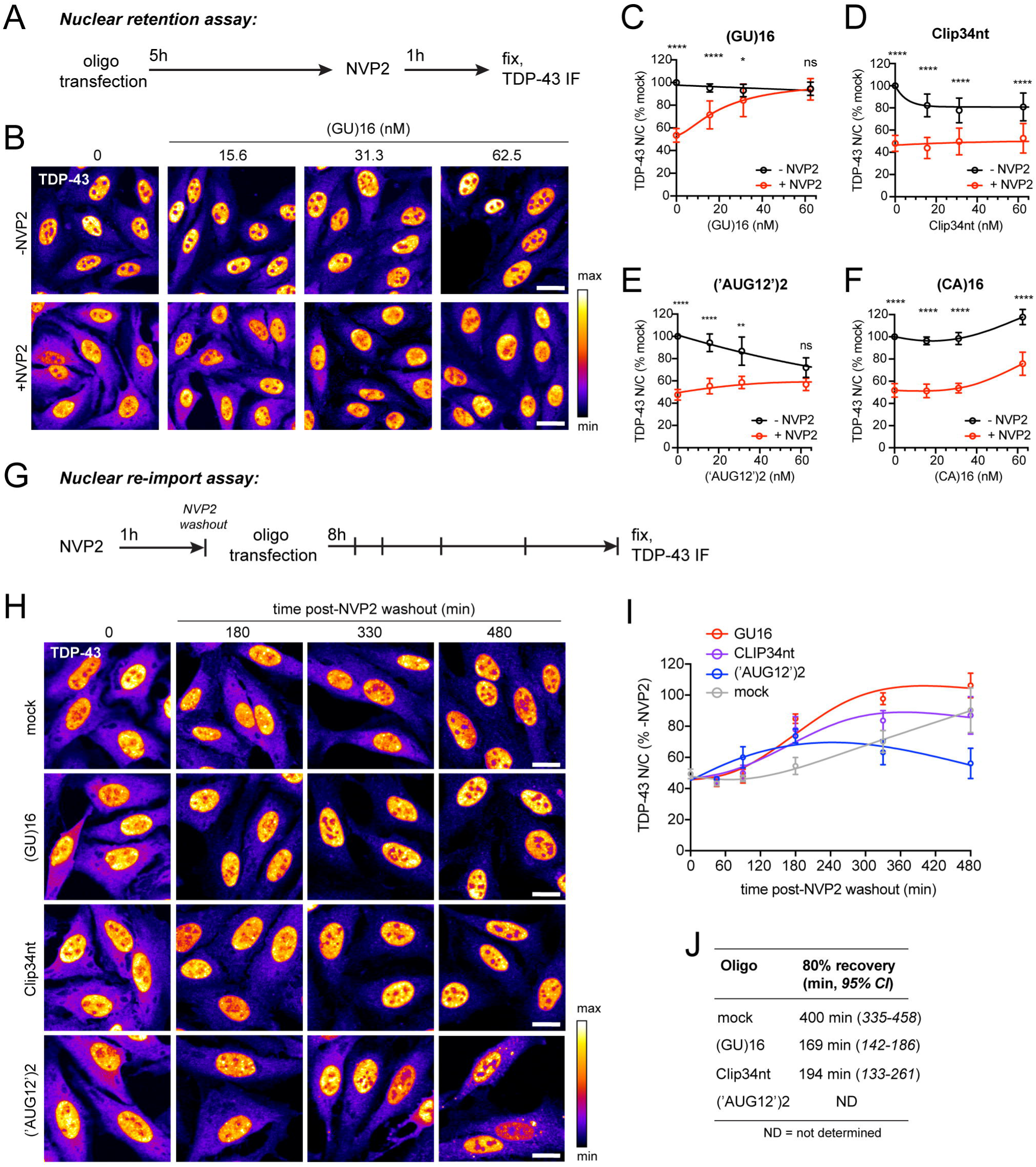
Multivalent GU-oligonucleotides attenuate transcriptional blockade-induced TDP-43 mislocalization and accelerate nuclear re-import. A. Schematic of the nuclear retention assay, in which oligonucleotides were first transfected into HeLa cells, incubated for 5 h, and TDP-43 mislocalization induced by NVP2 transcriptional blockade (250 nM for 1 h), prior to fixation and TDP-43 immunostaining. B. Representative TDP-43 immunofluorescence images, nuclear retention assay. Scale bar = 20 µm. C-F. TDP-43 N/C (% mock-transfected cells, i.e. Lipofectamine only) in (GU)16 (C), Clip34nt (D), (‘AUG12’)2 (E), or (CA)16 (F)-transfected cells with or without NVP2 treatment. Mean ± SD of ≥7 biological replicates. Note: -NVP2 curves are the same as in fig 2D. See also **figures S2-S3**. G. Schematic of the nuclear re-import assay, in which TDP-43 mislocalization was first induced by NVP2 treatment (250 nM for 1 h), followed by NVP2 washout and oligonucleotide transfection. A subset of cells were fixed immediately after NVP2 treatment and at timepoints up to 8 h for TDP-43 immunostaining. H. Representative TDP-43 immunofluorescence images, nuclear re-import assay. Scale bar = 20 µm. I. TDP-43 N/C normalized non-NVP2-treated cells at each timepoint. Mean ± SD of 3 biological replicates. J. Non-linear regression analysis of time to 80% recovery (minutes). 95% confidence intervals are shown. In B, H: the intensity histogram for each image was independently spread between the dimmest and brightest pixels and a pseudo-color linear LUT covering the full range of the data was applied (see legend). In C-F: NS= not significant, *p<0.05, **p<0.01, ***p<0.001, ****p<0.0001 by 2-way ANOVA with Tukey’s post- hoc test.

To investigate (GU)16-mediated attenuation of TDP-43 mislocalization from other perturbations, we tested the acute ablation of RanGAP1, the Ran GTPase-activating protein, which has been implicated in nucleocytoplasmic transport defects in mutant *C9ORF72* and TDP-43 aggregation models (**fig S3**).^39,40^ DLD1 cells containing an auxin-inducible degron (AID) integrated at the endogenous RanGAP1 locus and TIR1 ligase inserted at the C-terminus of RCC1 via a self-cleavable P2A sequence were generously provided by the Dasso lab.^41^ Auxin drives the rapid TIR1-dependent ubiquitination and degradation of RanGAP1 in this cell line within 2h (**fig S3A-B)**, accompanied by severe TDP-43 cytoplasmic mislocalization (**fig S3B**). Compared to mock-transfected cells, RanGAP1-AID cells transfected with (GU)16 for 5 h prior to auxin treatment showed dose-dependent attenuation of TDP-43 mislocalization (**fig S3C-D**). The effect was modest compared to NVP2, in part due to poor transfection efficiency of the DLD1 cells, which showed only ∼25-30% (GU)16 nuclear uptake compared to HeLa cells (**fig S3E**). Additionally, RanGAP1 ablation likely inhibits Ran-regulated nucleocytoplasmic transport, including TDP-43 nuclear re-import, which is not affected in the NVP2 assays.

Having observed that (GU)16 promotes TDP-43 nuclear retention when introduced prior to TDP-43 mislocalization, next we tested the ability of GU-oligonucleotides to promote recovery after TDP-43 mislocalization (fig 3G). We reasoned that the recovery of TDP-43 nuclear localization after a pulse of NVP2 likely depends on the resumption of nascent RNA synthesis to repopulate nuclear TDP-43 RNA binding sites, combined with continuous Ran- and importin-dependent nuclear import. Thus, the return of TDP-43 to the nucleus might be accelerated by the addition of exogenous, multivalent GU-RNAs. Cells were treated with NVP2 for 1 h to mislocalize TDP-43 to the cytoplasm, followed by NVP2 washout and oligonucleotide transfection. Cells were fixed/immunostained for TDP-43 at multiple timepoints up to 8 h post-transfection to analyze the recovery of nuclear TDP-43 localization. Biotinylated oligos rapidly entered transfected cell nuclei at similar rates (**fig S2B-C)**. Compared to mock-transfected cells, both (GU)16 and Clip34nt significantly accelerated the recovery of nuclear TDP-43 localization (time to 80% recovery [95% CI]: 169 [142-186] and 194 [133-261] min, versus 400 [335-458] min, respectively), whereas (‘AUG12’)2-transfected cells failed to recover and showed persistent cytoplasmic mislocalization (fig 3H-J). These findings suggest that even after depletion of endogenous nuclear pre-mRNAs, mislocalized TDP-43 that is reimported into the nucleus can be retained via binding to exogenous GU-oligos. Moreover, although substantial (GU)16 and Clip34nt is also found in the cytoplasm (fig 2A), any cytoplasmic oligo-TDP-43 binding does not inhibit the nuclear import of TDP-43, further supported by the lack of importin β in crosslinked RNA pulldown assays (**fig S4A**).

### GU-oligos induce the formation of heterogeneous high molecular weight RNP complexes

Next, we tested if transfected GU-oligonucleotides induce high molecular weight RNP complexes in cells, consistent with our biochemical data (**fig 1**) and the predicted mechanism of TDP-43 nuclear retention by multivalent binding. HeLa cells were transfected with biotinylated oligonucleotides. After 5 h, transfected cells were UV-irradiated to form covalent RNA-protein complexes and analyzed by SAV RNA-pulldowns (**fig 4a**). Only trace TDP-43 binding to (CA)16 was observed. As we previously reported, in (GU)6-transfected cells, SAV precipitated predominantly 43-45 kDa TDP-43 monomers,^28^ whereas in (GU)16-transfected cells, SAV- precipitated numerous higher molecular weight bands that were not multiples of 43 kD. Clip34nt and (‘AUG12’)2 produced similar findings, although at slightly shifted molecular weights, likely as a result of the different sizes of the crosslinked oligos. Thus, longer GU-oligos may induce the formation of heterogeneous TDP-43-containing RNP complexes that incorporate other nuclear proteins. Immunolabeling for other RNA binding proteins (RBPs) in (GU)16 transfected cells showed that the localization of MATR3, FUS, and hnRNPL at steady-state and after NVP2 treatment were unaffected by (GU)16 (fig 4F-H). However, (GU)16 partially attenuated the NVP2-induced nuclear efflux of hnRNPA1 and hnRNPA2/B1 (fig 4B-E). Neither hnRNPA1 nor hnRNPA2/B1 are predicted to bind GU-rich motifs,^42,43^ but both have been shown to interact with TDP-43^44^ suggesting that the (GU)16 response may be mediated by protein-protein interactions. Consistent with this, hnRNPA1 and hnRNPA2/B1 were not identified as direct (GU)16 binders in the SAV RNA-pulldowns, nor other RBPs, including MATR3, ELAVL1, ELAVL3, hnRNPC, SNRPA, or PSF (**suppl fig 4B**). hnRNPL bound (CA)16 but not (GU)16, consistent with its preferred binding to CA-repeat motifs^45^ and supporting the sensitivity and specificity of the pulldowns. FUS exhibited modest, monomeric binding to (GU)16, which is not expected from its consensus motif (CAGGACAGCCAG),^46^ and was insufficient to promote FUS nuclear retention (fig 4G). Like TDP-43, the nuclear export of FUS is independent of XPO1 and likely occurs by passive diffusion.^26^ We and others have found that FUS nuclear localization depends on the abundance of nuclear transcripts.^28,47^ Presumably, the reason for the lack of (GU)16-induced FUS nuclear retention is the low level and strictly monovalent binding, which is not expected to prevent the (GU)16-FUS complexes from exiting nuclei by passive diffusion through NPCs.^45^

**Figure 4.**
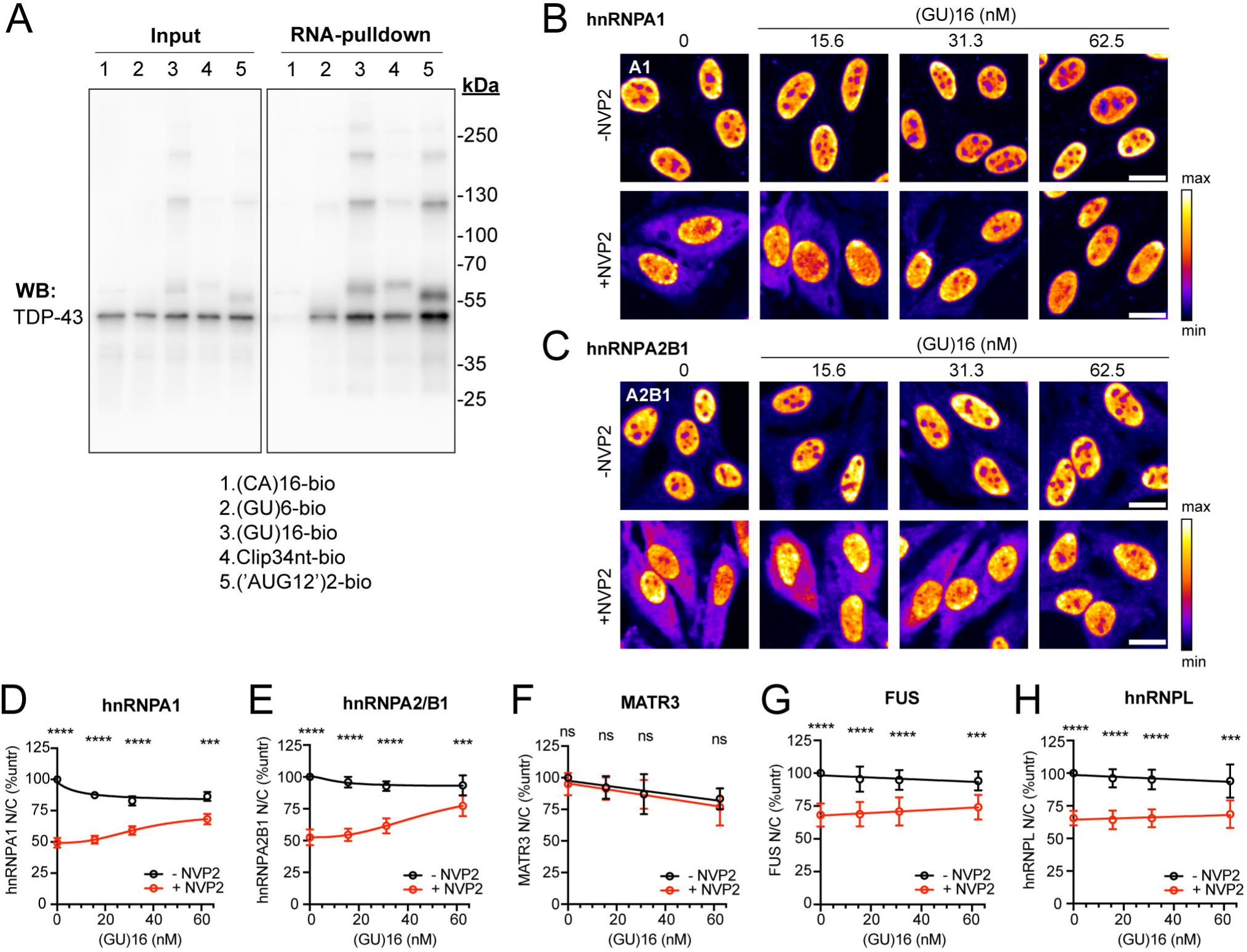
Formation of high-molecular weight oligo-TDP-43 complexes in transfected cells. A. SAV-RNA pulldowns in biotinylated oligo-transfected cells that were UV crosslinked before lysis and probed with rabbit anti-TDP-43. Results are representative of ≥3 biological replicates. See also **figure S4**. B-C. hnRNPA1 (B) and hnRNPA2B1 (C) immunofluorescence in cells transfected with (GU)16 for 5 h followed by 1 h transcriptional blockade with NVP2 (250 nM). The intensity histogram for each image was independently spread between the dimmest and brightest pixels and a pseudo-color linear LUT covering the full range of the data was applied (see legend). Scale bar = 20 µm. D-H. N/C ratio of hnRNPA1 (D), hnRNPA2B1 (E), Matrin-3 (F), FUS (G), and hnRNPL (H) in cells transfected with (GU)16 for 5 h followed by 1 h with or without NVP2 treatment. Mean ± SD of ≥4 biological replicates is shown. NS= not significant, ***p<0.001, ****p<0.0001 by 2-way ANOVA with Tukey’s post-hoc test.

### (GU)16-induced TDP-43 nuclear retention is RRM1,2-dependent

While effective in promoting RNase resistance, PS modification of RNA oligonucleotides has also been shown to increase protein binding.^48^ To verify that (GU)16-induced TDP-43 nuclear retention is mediated by specific binding to the TDP-43 RRM domains, we generated stable, monoclonal HeLa cell lines expressing V5-tagged wild-type TDP-43 (TDP43^WT^-V5) or TDP-43 containing five point mutations in the RRM1-2 domains that abolish RNA binding (TDP43^5FL^-V5)^49^ (fig 5A-D). As we previously reported, the steady-state N/C ratio of TDP43^5FL^-V5 was reduced compared to TDP43^WT^-V5 and TDP43^5FL^-V5 showed no NVP2-induced nuclear efflux.^28^ Next, the cell lines were transfected with (GU)16 for 5 h followed by NVP2 transcriptional blockade, and the localization of V5-tagged TDP-43 was analyzed by immunofluorescence. Like endogenous TDP-43 (**fig 3**), (GU)16 inhibited NVP2-induced TDP43^WT^-V5 mislocalization in a dose-dependent manner (fig 5A-B). However, (GU)16 had no effect on the localization of TDP43^5FL^-V5 (fig 5C-D), consistent with the notion that (GU)16 alters TDP-43 localization via specific binding to the RRM1,2 domain. Pertinent to the observation of (GU)16- induced alteration of TDP-43 nuclear granule morphology, TDP43-RRM mutant constructs have been reported to form nuclear bodies or anisosomes,^50–52^ which we also observed in transiently-transfected cells, though as we previously reported the frequency of cells containing TDP43-RRM mutant nuclear bodies is markedly higher in TDP43^5FL^-YFP expressing cells (∼100%) versus cells transiently expressing TDP43^5FL^-V5 (∼12%), suggesting a contribution of the YFP tag.^28^ The monoclonal TDP43^5FL^-V5 stable cell line generated here displays only rare nuclear bodies (<1% of cells, not shown), and thus is not suitable for analysis of potential effects of GU-oligo transfection on TDP43-RRM mutant nuclear bodies.

**Figure 5.**
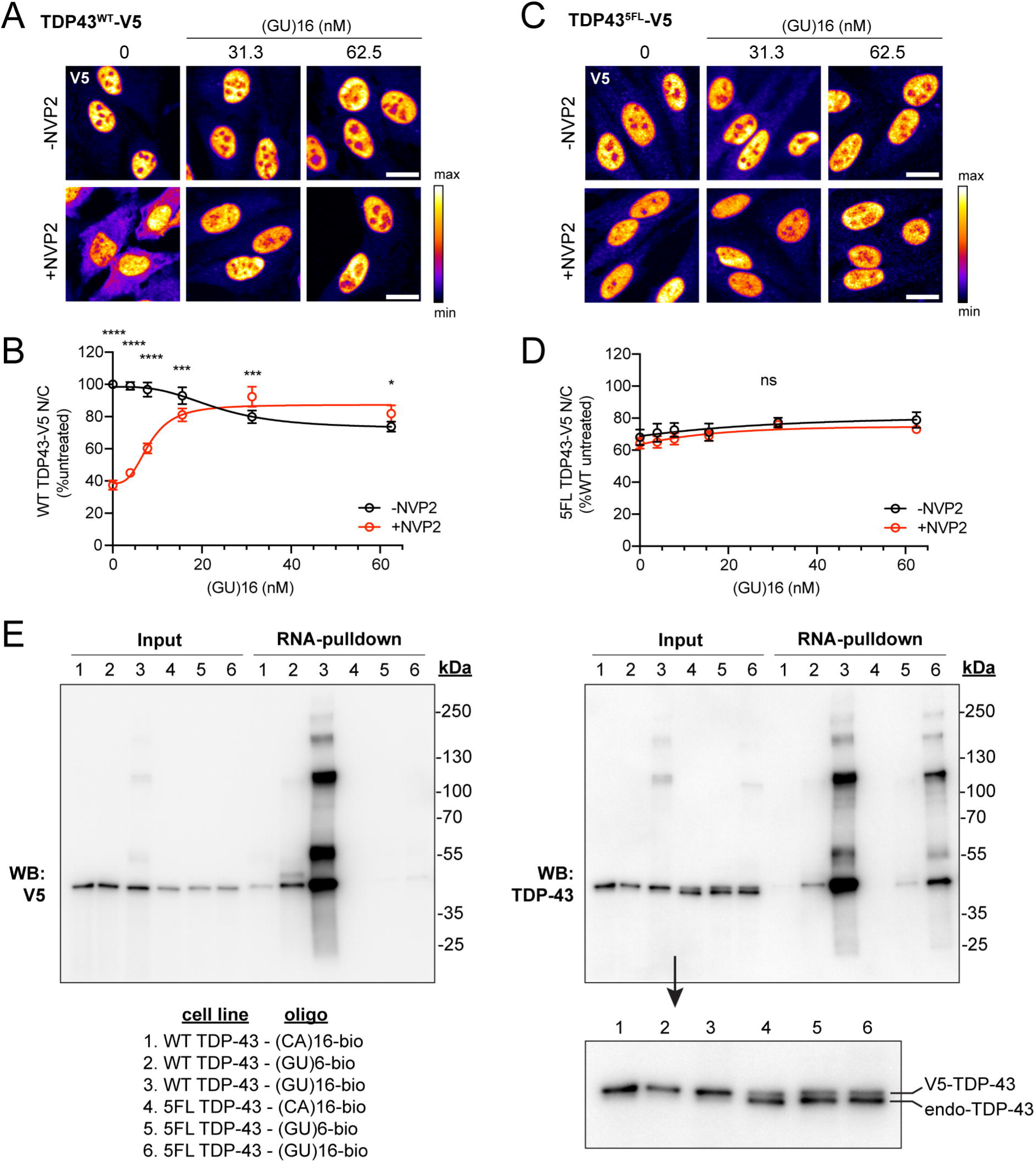
TDP-43 binding to (GU)16 is RRM1,2-dependent. A-B. V5 immunofluorescence (A) and N/C ratio (B) from TDP43^WT^-V5 monoclonal HeLa cells transfected with (GU)16 for 5 h, +/- subsequent 1 h NVP2 treatment. C-D. V5 immunofluorescence (C) and N/C ratio (D) from TDP43^5FL^-V5 monoclonal HeLa cells transfected with (GU)16 for 5 h, with or without subsequent NVP2 treatment. E. SAV-RNA pulldowns from biotinylated oligo-transfected cells that were UV crosslinked before lysis and probed with mouse anti-V5 (left) or rabbit anti-TDP-43 (right). Inset demonstrates ∼1.4 kDa separation between V5-tagged and endogenous TDP-43 and the expected presence (TDP43^WT^-V5 cells, lanes 1-3) or absence (TDP43^5FL^-V5 cells, lanes 4-6) of TDP-43 autoregulation. In A,C, the intensity histogram for each image was independently spread between the dimmest and brightest pixels and a pseudo-color linear LUT covering the full range of the data was applied (see legend). Scale bar = 20 µm. In B,D, N/C is expressed as % mock transfected cells. Mean ± SD of 4 biological replicates is shown. NS= not significant, *p<0.05, ***p<0.001, ****p<0.0001 by 2-way ANOVA with Tukey’s post-hoc test.

To further validate the direct (GU)16 interaction with RRM1-2, we performed SAV pulldowns from oligo- transfected TDP43^WT^-V5 or TDP43^5FL^-V5 cell lines (fig 5E). Immunoblots of the pulldowns developed with V5 antibodies (fig 5E**, left)** showed that transfection with (GU)6 and (GU)16 induced high molecular weight TDP43^WT^-V5 crosslinks, whose pattern was virtually identical to the crosslinks with endogenous TDP-43 (fig 5E**, right**), except for the small increase in size resulting from the 1.4kDa V5 tag. A barely detectable interaction of TDP43^5FL^-V5 with (GU)16 was noted (fig 5E**, left**). TDP43^WT^-V5 but not TDP43^5FL^-V5 showed trace binding to (CA)16). As previously demonstrated,^36^ TDP43^WT^-V5 but not TDP43^5FL^-V5 downregulated endogenous TDP-43 expression via autoregulation (fig 5E**, inset**).^36^ These data confirm that (GU)16 regulates TDP-43 localization by specific direct binding to the RRM1-2 domains.

### GU-repeat containing oligonucleotides cause TDP-43 loss of function

GU-rich oligonucleotides exhibit low nanomolar affinity for TDP-43^9,33–36^ and PS modification increases oligonucleotide protein binding.^48^ We suspected that although (GU)16 preserves steady-state TDP-43 nuclear localization (fig 2-3), high-affinity binding of PS-protected (GU)16 to TDP-43 may divert TDP-43 from endogenous RNAs and disrupt TDP-43 splicing regulatory function. To test this, we analyzed TDP-43 cryptic exon repression in *EPB41L4A*, a TDP-43-regulated transcript in HeLa cells^12,15^ and human neurons^53^ (**fig 6**). To assess *EPB41L4A* sensitivity as a TDP-43 functional readout, we first generated samples in which TDP-43 protein was depleted by approximately 50% (24 h post-siRNA transfection), versus cells in which TDP-43 was fully depleted by CRISPR (fig 6A).^54^ RT-PCR confirmed that *EPB41L4A* cryptic exon inclusion is detectable even after 50% loss of TDP-43 protein (fig 6B). Cells transfected with (CA)16 for 6, 18, or 24h showed no *EPB41L4A* cryptic exon inclusion, in contrast to (GU)16-transfected cells where the *EPB41L4A* cryptic exon was detectable by 6h and increased by 18h. To quantitatively compare TDP-43 loss of function (LOF) between oligonucleotides and account for the decrease in *EPB41L4A* mRNA with TDP-43 knockdown,^12^ we designed a qRT-PCR assay to quantify *EPB41L4A* cryptic exon versus mRNA levels, normalized to an internal spike-in control (fig 6C). Clip34nt, (‘AUG12’)2, and (GU)16 transfection all disrupted *EPB41L4A* cryptic exon repression, but interestingly the Clip34nt-induced LOF was much less severe than (‘AUG12’)2 and (GU)16. Similar results were obtained for a cryptic exon in *ARGHAP32* (**fig 6D**).^15^

**Figure 6.**
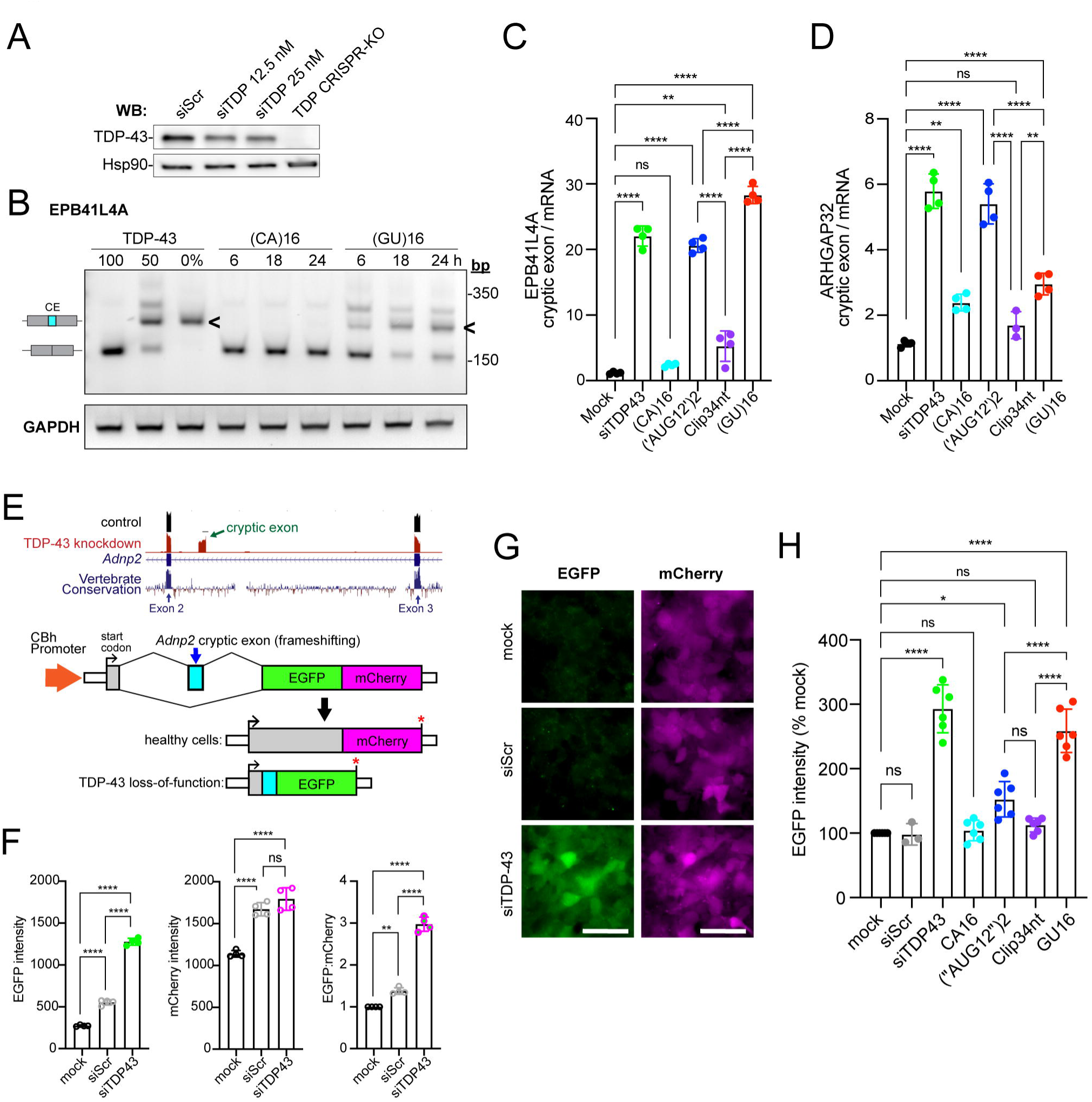
GU-repeat oligonucleotides disrupt TDP-43 function in cryptic exon repression. A. TDP-43 immunoblot 24 h-post transfection with scrambled siRNA (siScr) or 12.5-25 nM TDP-43 siRNA (siTDP), compared to a stable TDP-43 CRISPR knockout (KO) cell line. Hsp90 is used as a loading control. B. RT-PCR of *EPB41L4A* cryptic exon (CE <) inclusion in (CA)16 vs. (GU)16-transfected cells at 6, 18, and 24 h post-transfection. Controls for TDP-43 expression level include untreated cells (100%), TDP-43 siRNA-treated cells, 24 h (50%), and TDP-43 CRISPR knockout cells (0%). Representative of >3 biological replicates. For primers, see **table S2**. C. qRT-PCR of *EPB41L4A* cryptic exon inclusion in oligonucleotide-transfected cells, expressed as a ratio of cryptic exon to *EPB41L4A* mRNA. TDP-43 expression controls are as in (A-B). Mean ± SD of n=4 biological replicates. D. qRT-PCR of *ARHGAP32* cryptic exon inclusion in oligonucleotide-transfected cells, expressed as a ratio of cryptic exon to *ARHGAP32* mRNA. TDP-43 expression controls are as in (A-B). Mean ± SD of n=4 biological replicates. A single outlier was removed from Clip34nt (7.6) via Grubbs test (α=0.05). E. Schematic of bichromatic cryptic exon reporter, utilizing the TDP-43 cryptic exon from mouse *Adnp2*, in which TDP-43 knockdown induces a shift from mCherry to EGFP expression. F. EGFP intensity, mCherry intensity, and EGFP:mCherry ratio in a stable HEK293 reporter cell line that was mock transfected vs. scrambled siRNA (siScr) or TDP-43 siRNA (siTDP43) and imaged at 48 h. Mean ± SD of n=4 biological replicates. G. Representative widefield epifluorescence images of cells analyzed in F. The intensity histogram for all images was normalized to the min/max pixel intensity in the siTDP-43 condition. Scale bar = 50 µm. H. EGFP intensity of cryptic exon reporter cells 24h-post transfection with siScr, siTDP43, or the indicated oligonucleotides (62.5 nM). Mean ± SD of n=6 biological replicates. In C, D, F, H, ns = not significant, *p<0.05, **p<0.01, ****p<0.001 by one-way ANOVA with Tukey’s post-hoc test.

To evaluate these findings in an orthogonal system, we generated an HEK293 cell line stably expressing a bichromatic splicing reporter based on a TDP-43-regulated cryptic exon in mouse *Adnp2* (fig 6E). First, we validated that siRNA knockdown of TDP-43 induced a shift in fluorescence from mCherry to EGFP by live cell imaging (fig 6F-G). Indeed, compared to mock-transfected or scrambled siRNA (siScr)-transfected cells, siTDP-43 transfection induced a significant increase in EGFP intensity by 48h. At this timepoint, mCherry intensity had not yet decreased; rather, there was a modest non-specific increase in both siScr and siTDP-43- transfected cells, suggesting that the induction of EGFP expression is the most sensitive and specific marker of TDP-43 LOF in this cell line. Finally, we transfected the reporter cells with the RNA oligos and measured EGFP intensity at 24h (fig 6H). Consistent with data from endogenous cryptic exons, (‘AUG12’)2 and (GU)16 both induced a significant increase in EGFP intensity, whereas (CA)16 and Clip34nt had no effect. Together, these data suggest that GU oligo-induced TDP-43 LOF may depend on sequence, with pure GU-repeat-containing motifs causing greater LOF than A,C-interspersed, G,U-rich sequences. Further systematic testing is needed.

## DISCUSSION

We recently reported that TDP-43 is sequestered in the nucleus of healthy cells by binding to endogenous nuclear GU-rich RNAs, thus restricting its passive nuclear efflux.^28^ In that study, we found that competitive displacement of TDP-43 from endogenous nuclear RNAs by monovalent (GU)6 freed TDP-43 for passive diffusion from the nucleus. Here, we report that multivalent (GU)16 oligonucleotides sequester TDP-43 within the nucleus by forming high molecular weight RNP complexes that exceed the passive limits of the NPC. Transfection with low nanomolar (GU)16 prevented transcriptional blockade-induced TDP-43 mislocalization and attenuated TDP-43 mislocalization due to RanGAP1 ablation. Moreover, (GU)16 and Clip34nt accelerated TDP-43 re-import after cytoplasmic mislocalization. These data further support a reaction-diffusion model of RNA-based nuclear retention of TDP-43^28^ and provide proof of principle that exogenous GU-RNAs can promote TDP-43 nuclear localization. Disruption of TDP-43 function induced by oligonucleotides with pure GU-repeats was largely absent in cells transfected with interspersed Clip34nt, suggesting that future sequence optimization could lead to RNAs that support both TDP-43 nuclear localization and function.

### RNA sequence-specific drivers of multivalent TDP-43 RNPs

As previously reported,^32,55^ biochemical assays showed that longer GU-oligonucleotides promoted multivalent TDP-43 binding, compared to monovalent (GU)6 **(fig 1**). Both native EMSA assays and BS3 cross- linking-SDS-PAGE demonstrated stronger TDP-43 RRM1,2 multimerization for 34-nt Clip34nt and 36-nt (GU)16 compared with 24-nt (‘AUG12’)2. The RNA length-dependent formation of multivalent TDP-43-RNA complexes depends on RRM1-RRM2-mediated intramolecular cooperativity during binding to target RNA.^32^ Although binding affinity was not measured in the current study, past studies demonstrate a higher affinity for pure GU-repeats than interspersed sequences,^9,33–36^ which could also contribute to the increased multivalency that we observed for (GU)16. Indeed, a prior report found a 50-fold increase in binding affinity for Clip34nt- (GU)6 compared to Clip34nt,^34^ which lacks any GU tandem repeats. RNA secondary structure may also play a role, since Clip34nt is predicted to form a hairpin, while (GU)16 and (‘AUG12’)2 are unstructured. The potential effect of RNA secondary structure on TDP-43-RNA binding is yet unexplored.

### Oligonucleotide-induced regulation of TDP-43 localization in living cells

Upon oligonucleotide transfection into cells, additional factors likely contribute to the overall effect on TDP-43 binding and localization. Competition for oligo binding between TDP-43 and other RBPs with a similar motif preference presumably contributes to motif-specific differences in cells that is difficult to predict from biochemical assays, including differences in the localization of the oligonucleotides themselves (fig 2A-B). Surprisingly, oligo localization within cells did not predict the effect on TDP-43 nucleocytoplasmic localization and was inversely correlated, with (‘AUG12’)2 being the most nuclear and (GU)16 being equally distributed between the nucleus and cytoplasm. Future proteomic analysis could enable the identification of nuclear vs cytoplasmic oligo-bound proteins in transfected cells. The dose-dependency of oligo-induced effects (**fig 3 and fig S3**) suggests that the intracellular concentration of the oligos must reach a critical level, likely approaching the concentration of TDP-43, to alter its localization. Our results indicate that lipid-mediated transfection of synthetic oligos, in principle, meets such a requirement. Though only nanomolar concentrations were introduced, the transfection liposomes might intracellularly concentrate the oligos beyond what was added to the transfection mix, enabling successful oligo competition for nuclear TDP-43 binding.

TDP-43 LLPS regulates the native size of TDP-43 nuclear RNP complexes which in turn dictates TDP- 43 nucleocytoplasmic localization.^56^ Since GU-RNA modulates TDP-43 LLPS in a length- and motif-dependent

manner,^33,55^ the observed GU-oligo-induced effects on TDP-43 localization (and function, discussed below) may be influenced by changes in TDP-43 LLPS. RNA motifs containing pure GU repeats differ in their ability to promote TDP-43 phase transition compared to GU intermixed with A and C.^55^ Indeed, image analysis of oligo-induced changes in nuclear TDP-43 granules, likely composed of numerous TDP-43 RNPs, showed marked differences between oligos containing pure GU-repeats ((GU)16 and (‘AUG12’)2) versus the intermixed sequence of Clip34nt (**fig S1**). Namely, (‘AUG12’)2 and (GU)16 induced an increase in TDP-43 nuclear granule number while reducing mean granule size. We suspect that although both oligos induced the formation of multivalent, heterogeneous, TDP-43-containing RNP complexes (fig 4A), these RNPs are likely much smaller than TDP-43 RNPs involving endogenous RNAs,^31^ which induce cooperative TDP-43 binding and the formation of very large (>700-2000 kD) RNP complexes.^56,57^ Future studies utilizing size-exclusion chromatography could enable measurement of the native size of the GU-oligo induced granules for correlation with effects on TDP-43 localization.

Differential LLPS dynamics, together with binding affinity and RBP competition, may account for the variability we observed between (GU)16 and Clip34nt in the nuclear retention and nuclear reimport assays (**fig 3**). When introduced prior to transcriptional blockade (fig 3A-F), the oligos encounter a markedly different nuclear milieu than after NVP2 treatment (**fig 3G-J),** which rapidly depletes nuclear RNAs and induces the translocation of many RBPs to the cytoplasm.^28^ Whereas high affinity (GU)16 promoted TDP-43 localization in both paradigms, Clip34nt only showed efficacy in the re-import assay, suggesting that the nuclear RNA- and RBP-depleted conditions may enable more optimal Clip34nt-binding and, potentially, modulation of TDP-43 LLPS. Our findings are reminiscent of biochemical studies in which, unlike A(GU)18, Clip34nt-induced TDP-43 LLPS effects were blunted in cell lysates compared to purified assays.^33^ Future dedicated LLPS assays and cross-linking RNA pulldowns from the re-import assay could aid in determining if improved Clip34nt efficacy corresponds to more potent induction of high molecular weight TDP-43 RNP complexes in the re-import assay.

### TDP-43 functional disruption

The observation that transfected oligos disrupt TDP-43 function (**fig 6**) is predicted by the presumed mechanism of action, whereby GU-rich oligos competitively displace TDP-43 from endogenous pre-mRNA binding sites and promote the formation of ‘artificial’ RNP complexes. The high affinity, near-irreversible binding

of PS-modified GU-oligos to TDP-43 may be the primary driver of TDP-43 LOF, since the extent of cryptic exon derepression correlated with predicted oligo affinity ((GU)16 > (‘AUG12”)2 >> Clip34nt). However, other factors may contribute. For example, (‘AUG12’)2 also drives substantial TDP-43 cytoplasmic mislocalization, thus depleting the nuclear pool of TDP-43. As discussed above, shifts in TDP-43 nuclear granule morphology (**fig S1**) suggest potential oligo-induced modulation of TDP-43 LLPS which differs by oligo length and sequence. TDP-43 LLPS regulates its splicing function by promoting condensation at a subset of RNA regulatory sites characterized by specific lengths and RNA motifs.^31^ Thus, oligo-induced alterations in TDP-43 phase condensation may produce target-specific alterations in function across the transcriptome, which could be further analyzed by RNA sequencing. The finding that Clip34nt—an interspersed, non-GU-repeat sequence derived from a known endogenous TDP-43 binding site—largely spared TDP-43 nuclear function, suggests that screening similar endogenous motifs from CLIPseq datasets may yield additional oligonucleotide candidates to promote TDP-43 nuclear localization without perturbing its function.

### Limitations of the study

This study exclusively analyzed oligonucleotide transfection of dividing cell lines. Effects in dividing cells may not predict outcomes in neurons due to differences in endogenous RNA abundance, composition, and dynamics (e.g. synthesis, stability, modifications, and transport). Efforts are ongoing to overcome technical hurdles to studies in human neurons, including optimization of oligonucleotide length and base chemistry informed by antisense oligonucleotides, and the generation of endogenously expressed RNAs that could overcome constraints of length, delivery, and variability in expression that limit the potential for translation. Additionally, the current study is limited to acute (5-24 h) time periods and relies on severe perturbations of RNA synthesis and nucleocytoplasmic transport intended to target the molecular regulation of TDP-43 localization but of uncertain relevance to disease. Transitioning to human neurons will facilitate longer-term studies of the chronic effects of synthetic oligonucleotides on TDP-43 localization and function, including neuron-specific TDP-43 splicing regulatory targets (e.g. *STMN2,*^16^ *UNC13A,*^59^ NPTX2^58^) and other disease-relevant phenotypes, including TDP-43 solubility. Clip34nt was recently reported to disaggregate TDP-43 inclusions formed by optogenetic clustering in HEK293 cells and human neurons.^37,60^ Though TDP-43 solubility/disaggregation was not evaluated in the current study, together these findings highlight the potential

for RNA-based mechanisms to regulate TDP-43 localization, function, and solubility, which warrants systematic analyses going forward.

## Supporting information

Zhang et al. supplemental information

## ACKNOWLEDGEMENTS

We thank Fang Yang, Svetlana Vidensky, and Lyudmila Mamedova for expert technical assistance, and Angie Rubin and Barbara Smith for administrative support. Thank you to the Mary Dasso lab, including Vasilisa Aksenova, for generously providing the RanGAP1-AID cells and for helpful discussions. The graphical abstract was created with BioRender.com. This research was supported by NINDS/NIA R01NS123538, NINDS R03NS127011, the Muscular Dystrophy Association, the Guy McKhann Scholar Award (LH), and a JHU Catalyst Award (LH).

## AUTHOR CONTRIBUTIONS

Conceptualization PK, LH; Methodology XZ, TD, TC, VT, RC, JL, PK, LH; Investigation XZ, TD, PK, LH; Formal analysis XZ, TC, PK, LH; Visualization XZ, JL, PK, LH; Writing-original draft XZ, PK, LH; Writing-review & editing XZ, TD, JL, PK, LH; Supervision JL, PK, LH; Funding acquisition PK, LH.

## STAR METHODS

### RESOURCE AVAILABILITY

#### Lead contact

Further information and requests for resources and reagents should be directed to and will be fulfilled by the lead contact, Lindsey Hayes.

#### Materials availability

Plasmids and cell lines generated in this study will be shared by the lead contact upon completion of a material transfer agreement.

#### Data and code availability

- All data reported in this paper will be shared by the lead contact upon request. Original Western blot images are included in the supplement (**figure S5**).
- This paper does not report original code.
- Any additional information required to reanalyze the data reported in this paper is available from the lead contact upon request.

## EXPERIMENTAL MODEL DETAILS

### Cell lines

A single cell-derived HeLa cell clone (ATCC; originally female-derived)^61^ was maintained in OptiMEM (Gibco/ThermoFisher) with 4% FBS. Monoclonal HeLa cell lines stably expressing V5-tagged wild-type or RRM-mutant TDP-43 were generated by transient transfection using puromycin selection followed by limiting dilution cloning. A monoclonal TDP-43 CRISPR-knockout HeLa cell line was generously provided by Shawn Ferguson^54^ and maintained in DMEM (Gibco/ThermoFisher) with 10% FBS. U2OS cells (ATCC; originally female-derived) were maintained in DMEM/F12 with 10% FBS. DLD1-wildtype cells (ATCC; originally male-derived) and DLD1-RanGAP1-AID cells generously provided by Mary Dasso^41^ were maintained in DMEM (Gibco/ThermoFisher) with 10% FBS. The HTv3 TDP-43 cryptic exon reporter cell line was generated by stably transfecting HEK293 cells (ATCC) with the *Adnp2* bichromatic reporter plasmid. Cells positive for mCherry were then sorted by FACS and maintained in DMEM with 10% FBS. All cells were grown at 37° in humidified air containing 5% CO2, frequently refreshed from frozen stocks validated by STR profiling (ATCC) and verified to be mycoplasma negative (Genlantis).

## METHOD DETAILS

### Oligonucleotides

Synthetic RNA oligonucleotides were obtained from IDT. Sequences and modifications are listed in **table S1**. Desalted lyophilized oligonucleotides were reconstituted in sterile, RNAse- and DNAse-free water and single-use aliquots were stored at -80°C.

### Oligonucleotide transfections

Cells were plated 24 h prior to transfection in optical #1.5 glass- (CellVis: HeLa, U2OS) or plastic-bottom (Ibidi: DLD1, HEK293) 96-well plates at densities targeting 60-70% confluence at the time of transfection. As previously described,^28^ synthetic oligonucleotides were transfected using Lipofectamine RNAiMax (ThermoFisher). RNA oligomers were diluted in Opti-MEM and mixed 1:1 with diluted lipofectamine (1.5-3 µL per 25 µL Opti-MEM). After a 5 min incubation at room temperature, transfection mixes were added to culture media at 10-30 µL transfection mix per 100 µL media in 96-well plates or scaled proportionally for 6- and 12- well plates for protein and RNA extraction, respectively. The lipofectamine concentration and volume of transfection complex per well were optimized for each cell line utilizing biotinylated oligonucleotides to maximize dose-dependent nuclear entry, via SAV-647 labeling. The oligonucleotide concentrations shown represent the final concentration following dilution in culture media. Mock transfection controls were treated with lipofectamine alone. For transcriptional inhibition, cells were treated with 250 nM NVP2 (Tocris, stock in DMSO) for the times indicated in the figure legends. For auxin-induced RanGAP1 ablation, DLD1 cells were treated for 2 h with 0.5 mM 3-indoleacetic acid (Sigma, stock in ethanol). At the time points indicated, cells were fixed in 4% paraformaldehyde/PBS (Electron Microscopy Sciences) for 15 min.

### Immunofluorescence

Paraformaldehyde-fixed cells were rinsed once with PBS and then blocked and permeabilized with 10% normal goat serum (NGS, Vector Labs) and 0.1% Triton-X-100 in PBS for 15-30 min at room temperature. Primary antibodies were applied for 1 h at room temperature or overnight at 4°C in 10% NGS/PBS. Following two PBS rinses, AlexaFluor-labeled secondary antibodies (ThermoFisher) were applied for 1 h at room temperature in 5% NGS/PBS. To visualize biotinylated oligomers, streptavidin-AF647 (ThermoFisher) was added together with secondary antibody at 1:500. Cells were rinsed with PBS containing Hoechst 33342 and stored in 50% glycerol/PBS until imaging.

### Cryptic exon reporter assays

Reporter constructs were designed based on the TDP-43-regulated cryptic exon found in the mouse gene *Adnp2,* which is flanked by an extended GT repeat. Briefly, the region between exon 2 and exon 3 was PCR amplified from the mouse genome and inserted into a pSLED bichromatic reporter construct as previously described.^62^ HEK293 cells were transfected with the reporter plasmid to generate stably transfected cells. Cells positive for mCherry were FACS-sorted to generate the HTv3 cell line. Cells were validated by transfection with scrambled vs. TDP-43 siRNA (Dharmacon) using DharmaFECT 1 according to the manufacturer’s instructions. At 48 h, cells were transferred to prewarmed FluoroBrite DMEM (ThermoFisher) supplemented with 10% FBS and Hoechst 33342 for 20 min prior to live imaging. Subsequent oligonucleotide transfections were performed in parallel with siRNA controls as described above.

### RNA extraction and RT-PCR

Cells were rinsed in PBS and lysed in RNA lysis buffer (Zymo). Total RNA was isolated using the Quick RNA miniprep kit (Zymo) following the manufacturer’s protocol with DNase digestion. cDNA synthesis was performed with the high-capacity cDNA reverse transcription kit (ThermoFisher). RT-PCR for gel electrophoresis was performed using Platinum II Hot-Start Green PCR Master Mix (ThermoFisher), and products separated on 2% EX e-gels containing SYBR Gold II (Invitrogen). qRT-PCR using custom TaqMan primers/probes was performed with TaqMan Fast Advanced Master Mix on a QuantStudio 3 Real-Time PCR System. Prior to cDNA synthesis, samples were spiked with equal volumes of Taqman Universal RNA Spike-In RT control (ThermoFisher) which was utilized as an internal standard. Primer/probe sequences are provided in **table S2**.

### RRM1-2 cloning and recombinant protein production

The RRM1-2 domain of human TDP-43, including amino acids 102-269, was PCR-amplified from wt-TDP43- tdTomato-HA plasmid (Addgene 28205, a gift from Zuoshang Xu)^63^ and cloned in frame with the N-terminal 6- His-SUMO-TEV-site into the SspI site of pET 6His-SUMO-TEV LIC (1S) (Addgene 29659, a gift from Scott Gradia) using the HiFi Assembly kit (NEB). NEBuilder Assembly Tool 2.0 was used to design RRM1-2 primers with 21-bp overlap with the destination vector. The correct sequence of the resulting pET 6His-SUMO-TEV- RRM1-2 plasmid was verified by restriction digests and Sanger sequencing.

6His-SUMO-RRM1-2 protein production was performed in BL21-DE3 *E. coli* cells (Thermo), using LB media with 50µg/ml Kanamycin. Cells transformed with the pET 6His-SUMO-TEV-RRM1-2 plasmid were plated on an LB-agar plate and grown at 37°C overnight. The next day, a 250 mL starter culture was initiated with a single large colony and grown overnight on a shaker at 30°C, reaching OD_600nm_ =∼1.4. 100 mL of the starter was added to 1L media in 2.8L Fehrnbach baffle-free flask and incubated at 30°C, 225 rpm for 3.5 h, reaching OD_600nm_ =∼0.4. The culture was supplemented with 0.3mM IPTG and incubated at 30°C, 100 rpm for 3.5 h before cell harvesting and washing with 10mM imidazole in PBS (pH7.4), freezing in liquid nitrogen, and storage at -80°C. Protein purification started by quickly thawing the cells in ∼15 mL room-temperature 10mM imidazole with protease inhibitor cocktail (Roche) and freshly added 200µM PMSF. Cells were lysed by sonication on ice and lysates were clarified at 26500G, 40 min, 4°C. The clarified lysate was incubated on a rotator (50 min, 4°C) with 1 mL (packed volume) HIS-Select HF Nickel Affinity Gel (Sigma). The beads were collected in 15 mL Poly-Prep chromatography column (Bio-Rad) and washed with 15 mL 10mM imidazole before stepwise elution with 3ml each of PBS, pH7.4 containing increasing Imidazole concentrations (25- 1000mM). Aliquots of the eluates were separated on a 4-20% SDS-PAGE gel, and peak fractions (corresponding to 50-150mM imidazole) were identified by Coomassie staining. Pooled peak fractions were dialyzed overnight in PBS, clarified at 21000G, 4°C, 3 min, and protein concentration was measured before aliquoting, snap-freezing in liquid nitrogen and storage at -80°C.

### EMSAs

Oligonucleotides were diluted to 10 pmol in binding buffer (20 mM HEPES, pH 7.6 containing 25 mM KCl, 1 mM TCEP, and 2 mM MgCl_2_).^32^ A serial dilution of SUMO-RRM1,2 protein (0-40 pmol) was prepared, mixed 1:1 with oligonucleotide, and incubated at room temperature for 1 h. Samples were diluted in 5X TBE running buffer (Invitrogen/ThermoFisher) and run on 10% polyacrylamide gels (Invitrogen/ThermoFisher) at 200 V for 45 min on ice in pre-chilled 1X Ultrapure TBE Buffer (Fisher Scientific). Gels were rinsed in 1X TBE and immersed in SYBR Gold nucleic acid gel stain (1:10,000) (Invitrogen/ThermoFisher) for 30 min prior to fluorescent imaging on an ImageQuant LAS 4000 gel imaging system (GE). Band density was quantified using ImageJ.

### Biotinylated RNA-pulldowns

HeLa cells were transfected as described above in 6-well plates and crosslinked for RNA pulldowns as previously described.^28^ Briefly, 5 h post-transfection, cells were rinsed with DPBS and UV crosslinked (λ=254 nm, 1.5 J/cm^2^ on ice) before harvesting by trypsinization. Cells were washed twice with DPBS, resuspended in lysis buffer (1% NP40 in PBS, pH 7.4 with protease inhibitor cocktail (Roche), and 2.5% v/v RNase inhibitor), and sonicated 10s on ice. After clarification of the lysates by centrifugation (21,000g, 5 min, 4°C) and protein concentration measurement, samples were diluted with lysis buffer to equal concentration (2-4 µg/µL). An equal total protein amount of lysate per sample was then added to DPBS-washed magnetic SAV-conjugated beads (Thermo-Fisher MyOneC1, ∼1 µL beads/ 15-20µg sample) and rotated at 4°C for 1.5 h. Beads were collected on a magnetic stand, supernatants removed and beads washed 5 times with ice-cold RIPA buffer (10mM Tris-HCl, pH 8.0, 5mM EDTA,1% Triton X-100, 0.5% Sodium Deoxycholate, 0.1% SDS, 150mM NaCl), transferred to the new set of tubes, rinsed twice with water and resuspended in 20 µL Laemmli sample buffer (Bio-Rad, 2X with 5% β-mercaptoethanol). After boiling for 5 min at 100°C, tubes were briefly centrifuged, placed on a magnetic stand, and the eluates separated by SDS-PAGE, along with aliquots of inputs prepared in Laemmli sample buffer. After transfer to PVDF membranes, immunoblots were blocked for 30-60 min with 5% non-fat dried milk (Biorad) in TTBS (0.05% Tween-20, TBS, pH7.4) and incubated overnight at 4°C with primary antibodies. After incubation with horseradish peroxidase-coupled secondary antibodies (20 min, room temperature) and washes with TTBS, ECL images were photographed using ImageQuant LAS400.

## QUANTIFICATION AND STATISTICAL ANALYSIS

### Microscopy and automated image analysis

Automated cell imaging was performed on an ImageXpress Micro Confocal high content microscope (Molecular Devices) as previously described.^28,64^ Fixed, immunostained cells were imaged with a 20x objective in spinning disc confocal mode with 60 µM pinhole at sub-saturating exposures. A 3 x 3 grid of nine non-overlapping images were routinely collected from each well yielding an average of 1000-3000 cells per well, depending on cell type and density. 96-well plate designs routinely incorporated 2-3 wells per dose or condition to serve as technical replicates, which were combined to generate the mean ± SD for each biological replicate shown in the figures. Background-corrected mean nuclear and cytoplasmic intensities and the nuclear/cytoplasmic (N/C ratio) were analyzed using the MetaXpress translocation-enhanced module (Molecular Devices). Nuclear TDP-43 granules were analyzed using a custom module set to detect nuclear granules ≥ 1 pixel (0.5 µm) in size and ≥4000 relative fluorescence units brighter than local background, among non-mitotic cell nuclei. All image analysis was performed on raw, unaltered images and uniformly filtered to exclude errors of cell identification (probe intensity = 0) or non-physiologic results (N/C ratio <0.1 or >100). Live cell imaging of HTv3 reporter cells was done in 20x widefield mode, and dual GFP/mCherry intensities analyzed with the MetaXpress multiwavelength translocation module. High resolution images of oligonucleotide localization and nuclear TDP-43 granules were obtained using a Zeiss LSM980 confocal microscope with a Plan-Apochromat 63x/1.4 oil objective, in confocal and Airyscan mode, respectively. For quantification of TDP-43 nuclear granules in Image J, individual nuclei were outlined, and surrounding pixels discarded (‘clear outside’ tool). Images were thresholded to the top 5^th^ percentile and the ‘analyze particles’ tool was used with lower limit ≥ 1 pixel (0.035 µm).

### Image processing for figures

Minimal processing was done using Adobe Photoshop 2024, limited to cropping and spreading the intensity histogram between the dimmest and brightest pixels in each image. In selected figures, the ‘fire’ pseudo-color look-up-table (LUT/ImageJ) was applied to aid in data visualization. The LUT is a quantitative and linear map that covers the full range of the data. All adjustments were linear (no gamma changes) and applied equally to the entire image. Immunoblots and gels were cropped for space and the intensity histogram was spread between the dimmest and brightest pixels. Unmodified gel images are shown in **figure S5**.

### Statistical analysis

Curve fitting and statistical analyses were performed using Prism software (Graphpad). Dose-response curves were fit by simple linear or non-linear regression analysis. Differences among groups were analyzed by ANOVA with post-hoc tests for multiple comparisons. The number of biological replicates (independent cell passages and experiments) was used as the N for all analyses, unless otherwise noted in the figure legend. Outliers were detected with the Grubbs (extreme Studentized deviate) method to detect a single outlier (α = 0.05). This resulted in one outlier being removed (**Fig 6D**).

