## Supplementary material for "Multivalent GU-rich oligonucleotides sequester TDP-43 in the nucleus by inducing high molecular weight RNP complexes": Zhang et al. supplemental information

### Figure S1

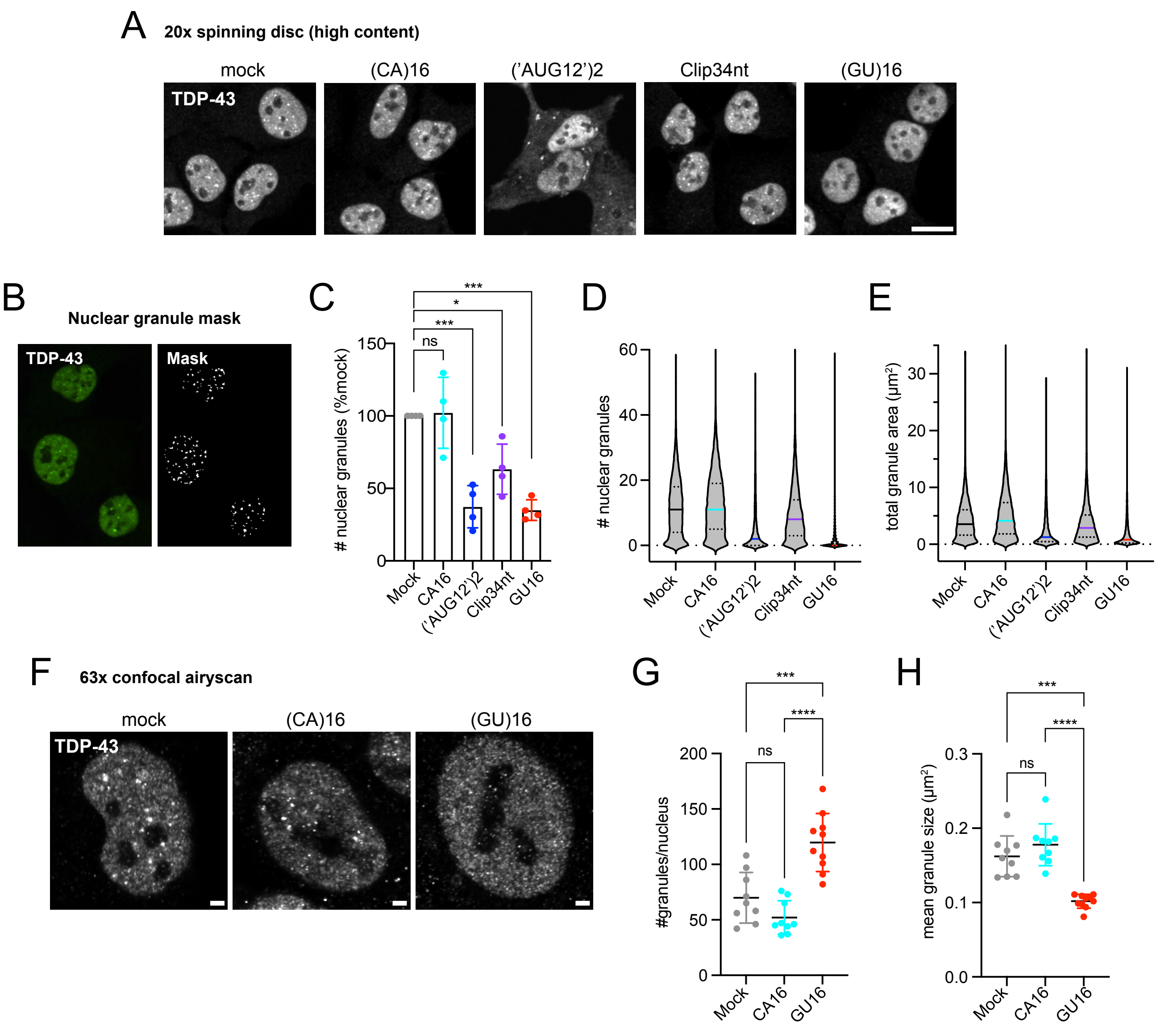


**Figure S1. GU-oligonucleotide transfection alters nuclear TDP-43 granule morphology (related to figure 2).** Oligo-transfected U2OS cells (125 nM, 24 h) were fixed and immunostained for TDP-43, and imaging/analysis of nuclear granule morphology was performed with 20x spinning disc high content microscope utilizing MetaXpress (**A-E**) or 63x confocal airyscan utilizing Image J (**F-H**). **A.** Representative 20x images. Scale bar = 20 μm. **B.** Representative nuclear granule mask generated with custom MetaXpress analysis module (lower limit 0.5 μm). **C.** Number of nuclear TDP-43 granules expressed as % mock-transfected cells. Mean ± SD of N=4 biological replicates. NS = not significant, *p<0.05, ***p<0.001 by one-way ANOVA with Tukey's post-hoc test. **D-E.** Violin plots of granule number (D) and total area/cell (E) from a representative replicate within C (n~4,000-12,000 cells per condition). Median, 25$^{th}$, and 75$^{th}$ percentiles are indicated. **F.** Representative 63x confocal airyscan images. Scale bar = 2 μm. **G.** Mean number of TDP-43 granules/nucleus quantified by Image J 'analyze particles' function (lower limit 0.035 μm). **H.** Mean TDP-43 granule size (μm). In G-H, N=9-10 cells/group. NS = not significant, ***p<0.001, ****p<0.0001 by one-way ANOVA with Tukey's post-hoc test.

### Figure S2

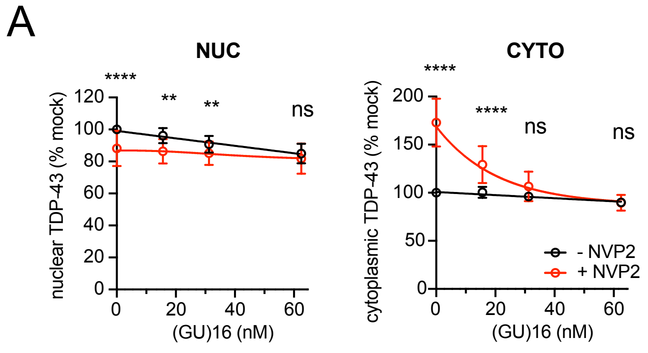


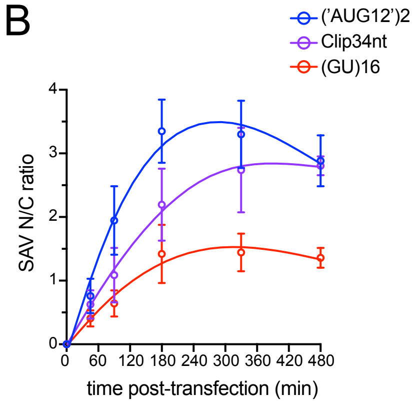


C

| Oligo | $t_{1/2}$ (min, *95% CI*) |
|---|---|
| ('AUG12')2 | 58 min (37-101) |
| Clip34nt | 106 min (60-268) |
| (GU)16 | 72 min (40-171) |

**Figure S2**. **Extended analysis, GU-oligo-induced nuclear TDP-43 retention (related to figure 3). A**. Nuclear and cytoplasmic intensity in (GU)16-transfected cells (% mock-transfected cells), with or without NVP2 treatment. These are the source data used to calculate the N/C ratios in fig 3C. Mean ± SD of n=15 biological replicates. NS= not significant, **p<0.01, ****p<0.0001 by 2-way ANOVA with Tukey's post-hoc test. **B.** SAV647 nuclear intensity in oligo-transfected cells during the NVP2 recovery assay in fig 3H-I. Mean ± SD from n=3 biological replicates. **C.** $t_{1/2}$ (min) for nuclear uptake calculated from (B) by non-linear regression. 95% CI is shown.

### Figure S3

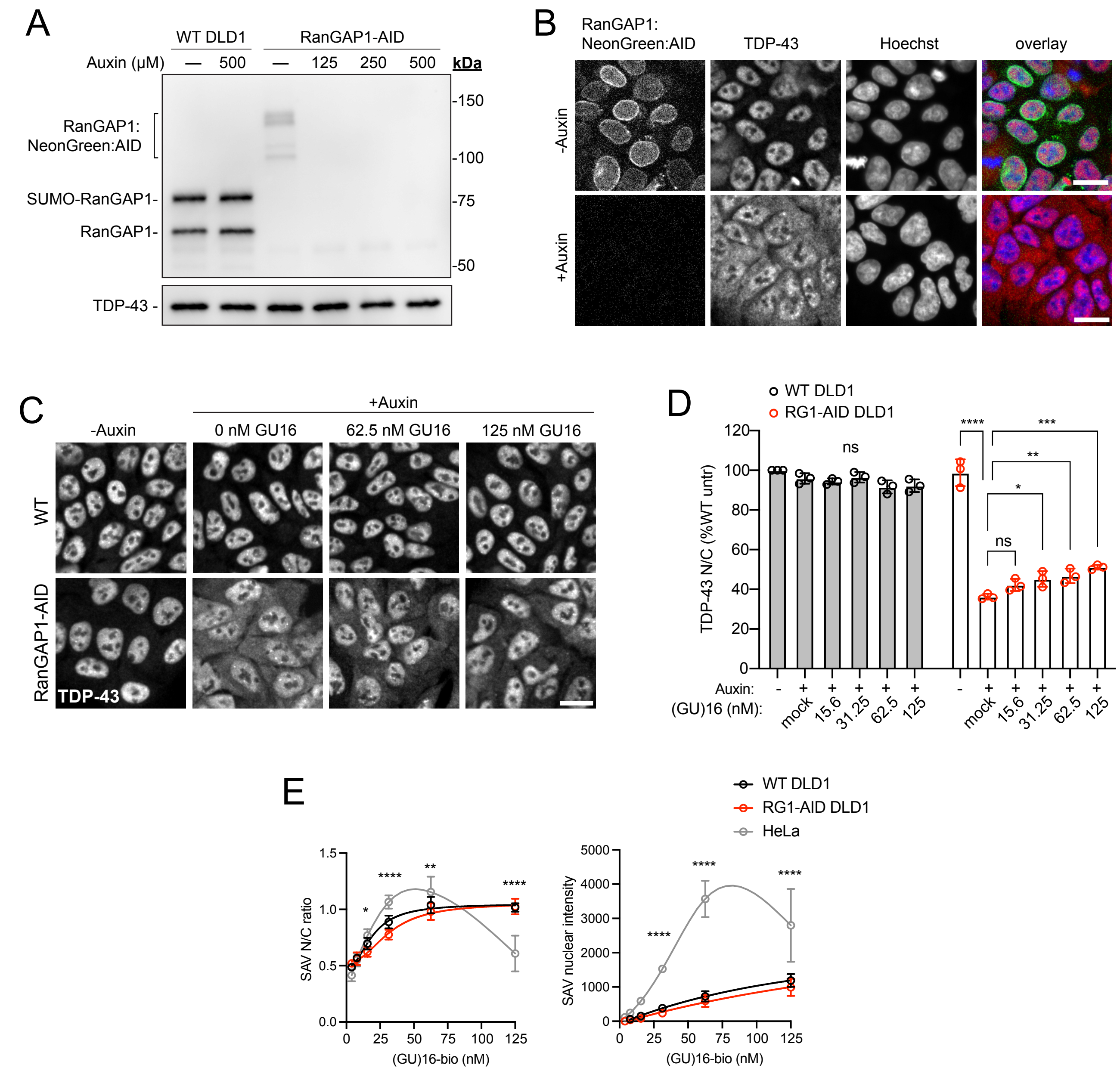


**Figure S3. (GU)16 attenuates TDP-43 mislocalization in RanGAP1-auxin inducible degron (AID) cells (related to figure 3)**. **A.** RanGAP1 immunoblot in wild-type (WT) and RanGAP1-AID cells ± 2 h auxin (indoleacetic acid) treatment. Distinct bands were seen as expected for RanGAP1 and SUMOylated RanGAP1. Note: the RanGAP1 AID cell line has reduced RanGAP1 expression (bracket) suggesting accelerated RanGAP1 turnover independent of auxin (perhaps due to excess TIR1). TDP-43 expression is unchanged. **B.** Loss of RanGAP1:NeonGreen:AID epifluorescence in RanGAP1 AID cells after 2 h 500 µM auxin treatment is accompanied by cytoplasmic mislocalization of TDP-43 (immunostained with AF568-labeled secondary). Scale bar = 20 µm. **C.** TDP-43 immunostaining in WT versus RanGAP1 AID cells transfected with increasing concentrations of (GU)16 for 5 h followed by 2 h 500 µM auxin treatment. Scale bar = 20 µm. **D.** TDP-43 N/C expressed as %WT untreated cells. Mean ± SD from n=3 biological replicates. NS= not significant, *p<0.05, **p<0.01, ****p<0.0001 by 2-way ANOVA with Tukey's post-hoc test. **E.** SAV647 N/C ratio (left) and nuclear intensity (right) in biotinylated (GU)16-transfected DLD1 vs. HeLa cells under optimized conditions for each cell type. Mean ± SD for ≥3 biological replicates. *p<0.05, **p<0.01, ****p<0.001 by 2-way ANOVA with Tukey's post-hoc test (all groups were compared; asterisks are shown for HeLa vs. RG1-AID comparisons only).

### Figure S4

A

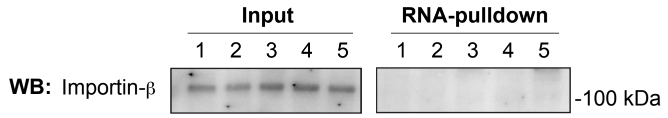


1.(CA)16-bio
2.(GU)6-bio
3.(GU)16-bio
4.Clip34nt-bio
5.('AUG12')2-bio

B

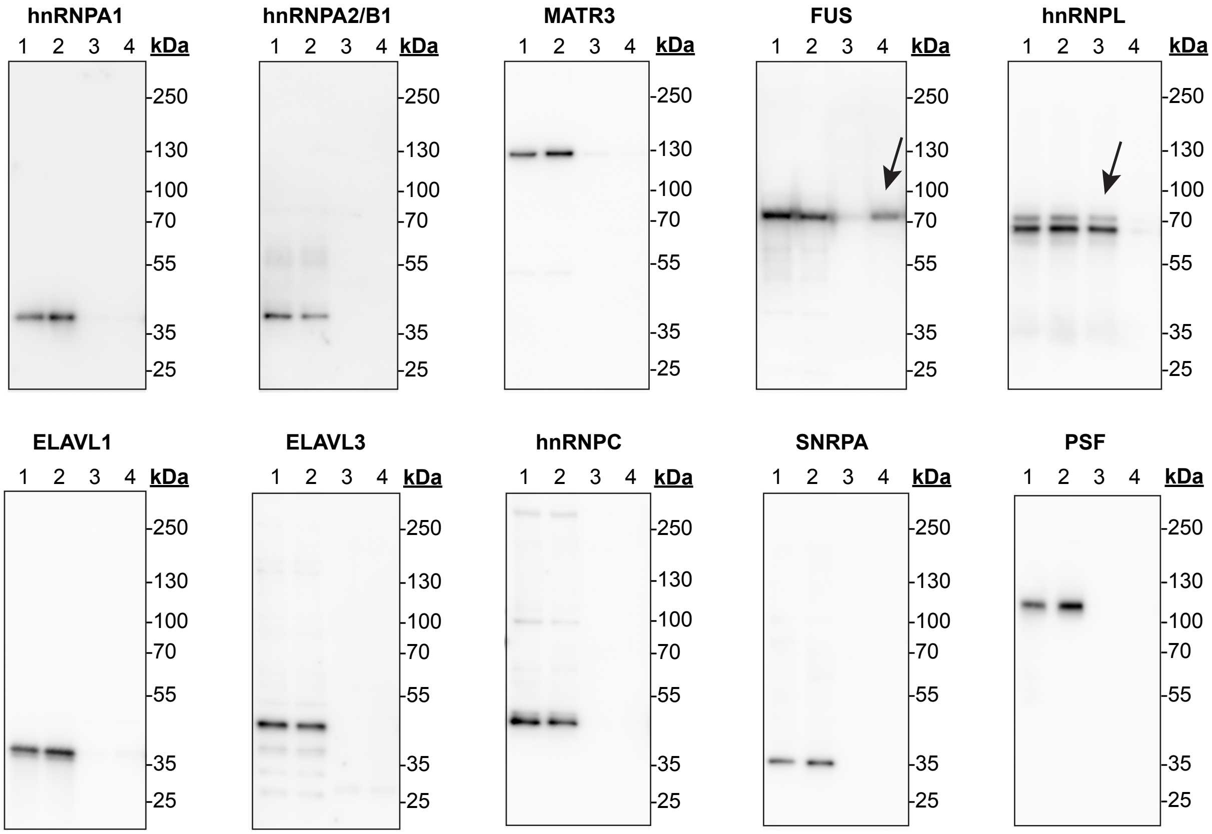


1. Input - (CA)16-bio
2. Input - (GU)16-bio
3. Pulldown - (CA)16-bio
4. Pulldown - (GU)16-bio

**Figure S4. SAV-RNA pulldowns probed for additional binding partners (related to figure 4).** SAV-RNA pulldowns from biotinylated oligo-transfected cells (62.5 nM, 5 h) that were UV crosslinked before lysis and probed with the indicated antibodies, to importin β (**A**), or a panel of nuclear RNA binding proteins (**B**). Note: the blot in A is a reprobe of the blot in fig 4a. Arrows in B indicate FUS binding to (GU)16 and hnRNPL binding to (CA)16.

### Figure S5

**Fig 1b**

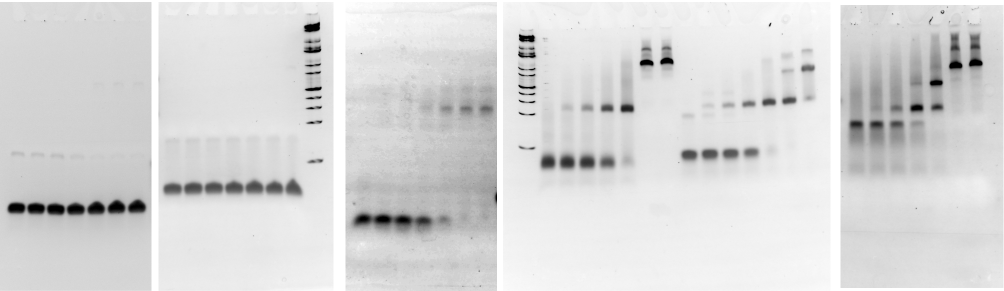

**Fig 1d**

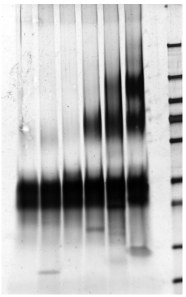

**Fig 4a**

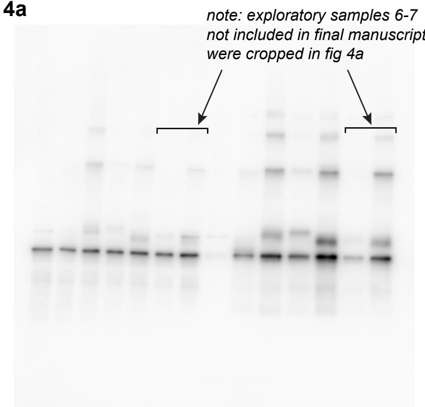


**Fig 5e**

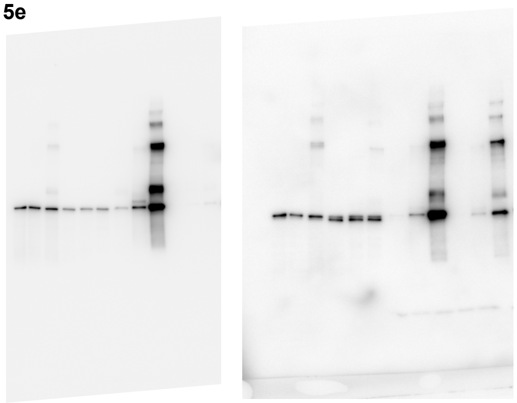

**Fig 6a**

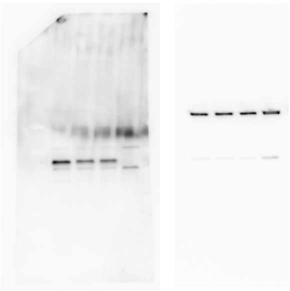

**Fig 6b**

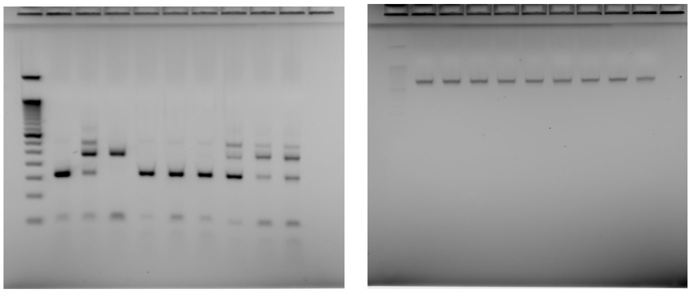

**Figure S5. Unmodified gel images, related to figures 1, 4, 5, and 6.**

#### SUPPLEMENTAL TABLES

##### Table S1. RNA oligonucleotides

| Oligo | RNA sequence with modifications<br>m: 2' O-methyl<br>*: phosphorothioate bond | ID # |
|---|---|---|
| GU6-biotin | 5'-mG*mU*mG* mU*mG*mU*mG*mU*mG* mU*mG*rU/3Bio/-3' | R78 |
| GU16-biotin | 5'-mG*mU*mG*mU*mG*mU*mG*mU*mG* mU*mG*mU*mG*mU*mG* mU*mG*mU* mG*mU*mG*mU*mG*mU*mG*mU*mG*mU*mG*mU*mG*rU/3Bio/-3' | R81, R102, R105 |
| CA6-biotin | 5'-mC*mA*mC*mA*mC*mA*mC*mA*mC*mA*mC*rA/3Bio/-3' | R75 |
| CA16-biotin | 5'-mC*mA*mC*mA*mC*mA*mC*mA*mC*mA*mC*mA*mC*mA*mC*mA*mC*mA* mC*mA*mC*mA*mC*mA*mC*mA*mC*mA*mC*mA*mC*rA/3Bio/-3' | R83, R107 |
| ('AUG12')2-biotin | 5'-mG*mU*mG*mU*mG*mA*mA*mU*mG*mA*mA*mU*mG*mU*mG* mU*mG*mA*mA*mU*mG*mA*mA*rU/3Bio/-3' | R95 |
| Clip34nt - biotin | 5'-mG*mA*mG*mA*mG*mA*mG*mC*mG*mC*mG*mU*mG*mC*mA*mG*mA*mG* mA*mC*mU*mU*mG*mG*mU*mG*mG*mU*mG*mC*mA*mU*mA*rA/3Bio/-3' | R76, R103 |
| A12-biotin | 5'-mA*mA*mA*mA*mA*mA*mA*mA*mA*mA*mA*rA/3Bio/-3' | R74 |
| A32-biotin | 5'-mA*mA*mA*mA*mA*mA*mA*mA*mA*mA*mA*mA*mA*mA*mA*mA*mA*mA* mA*mA*mA*mA*mA*mA*mA*mA*mA*mA*mA*mA*mA*rA/3Bio/-3' | R96 |

##### Table S2. Primers

| Target | Sequence | Source or catalog # |
|---|---|---|
| RT-PCR | | |
| EPB41L4A-F | GGACCTCCATATACTTTGTATTTTGGT | Tan *et al*., 2016 |
| EPB41L4A-R | AGCTGAGCAGCAGTGTTGAC | Tan *et al*., 2016 |
| GAPDH-F | ACCACAGTCCATGCCATCAC | IDT |
| GAPDH-R | TCCACCACCCTGTTGCTGTA | IDT |
| qRT-PCR | | |
| EPB41L4A cryptic exon | F - ACATATGCACACACACTCTCACA<br>Probe – CACTTCAGCCTGTCCTTT (FAM)<br>R - AGTCACCTTACAAAACAGAAGTCACA | ThermoFisher custom assay APRWNHN |
| ARHGAP32 cryptic exon | F - CAGCCAAATGCACAGCGAAT<br>Probe – AATGACCCAGTCAAAATA (FAM)<br>R - GGAGGAAGCATTTTGGAGGTTTCTA | ThermoFisher custom assay APT2G3K |
